# Advancing terrestrial biodiversity monitoring with satellite remote sensing in the context of the Kunming-Montreal global biodiversity framework

**DOI:** 10.1101/2022.04.25.489356

**Authors:** Joris Timmermans, W. Daniel Kissling

## Abstract

Satellite remote sensing (SRS) provides huge potential for tracking progress towards conservation targets and goals, but SRS products need to be tailored towards the requirements of ecological users and policymakers. In this *viewpoint* article, we propose to advance SRS products with a terrestrial biodiversity focus for tracking the goals and targets of the Kunming-Montreal global biodiversity framework (GBF). Of 371 GBF biodiversity indicators, we identified 58 unique indicators for tracking the state of terrestrial biodiversity, spanning 2 goals and 8 targets. Thirty-six shared enough information to analyse their underlying workflows and spatial information products. We used the concept of Essential Biodiversity Variables (EBV) to connect spatial information products to different dimensions of biodiversity (e.g. species populations, species traits, and ecosystem structure), and then counted EBV usage across GBF goals and targets. Combined with published scores on feasibility, accuracy, and immaturity of SRS products, we identified a priority list of terrestrial SRS products representing opportunities for scientific development in the next decade. From this list, we suggest two key directions for advancing SRS products and workflows in the GBF context using current instruments and technologies. First, existing terrestrial ecosystem distributions and live cover fraction SRS products (of above-ground biomass, ecosystem fragmentation, ecosystem structural variance, fraction of vegetation cover, plant area index profile, and land cover) need to be refined using a co-design approach to achieve harmonized ecosystem taxonomies, reference states and improved thematic detail. Second, new SRS products related to plant physiology and primary productivity (e.g. leaf area index, chlorophyll content & flux, foliar N/P/K content, and carbon cycle) need to be developed to better estimate plant functional traits, especially with deep learning techniques, radiative transfer models and multi-sensor frameworks. Advancements along these two routes could greatly improve the tracking of GBF target 2 (‘improve connectivity of priority terrestrial ecosystems), target 3 (‘ensure management of protected areas’), target 6 (‘control the introduction and impact of invasive alien species’), target 8 (‘minimize impact of climate change on biodiversity’), target 10 (‘increase sustainable productivity of agricultural and forested ecosystems’) and target 12 (‘increase public urban green/blue spaces’). Such improvements can have secondary benefits for other EBVs, e.g. as predictor variables for modelling species distributions and population abundances (i.e. data that are required in several GBF indicators). We hope that our *viewpoint* stimulates the advancement of biodiversity monitoring from space and a stronger collaboration among ecologists, SRS scientists and policy experts.

## 1 Introduction

Global biodiversity is rapidly declining and human impacts impair the distribution and abundance of wild species, the distinctness of ecological communities, and the extent and integrity of ecosystems (Díaz et al., 2019). Since the 1992 Rio Earth Summit, the Convention on Biological Diversity (CBD) has aimed to promote sustainable development and conservation of biodiversity through national policies. In December 2022, governments have released the final text of the Kunming-Montreal Global Biodiversity Framework (GBF) with a set of ambitious long-term goals (to be achieved in 2050) and specific targets (for 2030) that require immediate action (CBD, 2022a). Within the monitoring framework of this GBF, 371 headline, component and complementary biodiversity indicators —created by non-governmental organisations, research institutes and other partners of the Biodiversity Indicators Partnership (BIP)— have been identified to track progress towards achieving these goals and targets (CBD, 2022b). We believe that satellite remote sensing (SRS) could play a vital role in providing data to feed into these biodiversity indicators if ecological and policy requirements can be met. With this *viewpoint* article, we review the potential importance of SRS observations from current instruments and technologies for monitoring the state of terrestrial biodiversity in the GBF context and suggest key directions for advancing and developing SRS products and workflows.

Assessments of progress towards global conservation goals and targets require a multitude of datasets that can feed into biodiversity indicators. Besides *in-situ* monitoring, SRS has been suggested to play a vital role in the development of reliable indicators, especially in the terrestrial realm (Cavender-Bares et al., 2022; Pettorelli et al., 2016; Skidmore et al., 2015; Turner, 2014; Wang and Gamon, 2019). One advantage of SRS is that it acquires spatially contiguous data on biodiversity and ecosystems with a (near-)global extent, and with a high consistency over the lifetime of the mission. Nevertheless, within ecology and biodiversity science the use of SRS products has been mostly seen in the context of predictor variables to extrapolate in-situ observations of species distributions and population abundance into continuous variables in space and time (Pereira et al., 2017; Randin et al., 2020). This contrasts with the view that SRS observations can be used to monitor specific aspects of biodiversity directly from space (Alleaume et al., 2018; Pettorelli et al., 2016; Skidmore et al., 2021, 2015). Recent studies do indeed show that SRS can provide observations (Koskikala et al., 2020) and derived products (e.g. fractional cover, forest cover or land cover) for monitoring the distribution, fragmentation and heterogeneity of ecosystems (Lock et al., 2022). Moreover, with increasing amounts of new SRS products (Kuenzer et al., 2014) and new satellite missions emerging (Briottet et al., 2022; Cawse-Nicholson et al., 2021), the range of observations that can directly contribute to biodiversity indicators will further broaden. Examples include imaging spectrometers for estimating phylogenetic and trait diversity of plants (Asner et al., 2017; Helfenstein et al., 2022; Schneider et al., 2017), radar for mapping forest biodiversity (Bae et al., 2019), LiDAR sensors for measuring the 3D structure of ecosystems (Valbuena et al., 2020), or a combination of multispectral images from different satellite sensors to count large terrestrial mammals (Wu et al., 2023). However, an ecological perspective of how SRS can be improved in the context of the final Kunming-Montreal GBF is currently lacking, apart from a preliminary perspective on the 1 draft version (Cavender-Bares et al., 2022).

Requirements for developing new SRS products can be defined by assessing the feasibility, accuracy and (im)maturity of SRS products (Skidmore et al., 2021), e.g. regarding the potential of novel approaches (Skidmore et al., 2021, 2015), the validity of data products (Mayr et al., 2019) or the capability of processing chains (Paganini et al., 2016). Such assessments, often conducted by the remote sensing community, provide a high level of technological detail which is useful for space agency engineers and SRS scientists, but difficult to understand for ecologists and policy makers (Kuenzer et al., 2014; O’Connor et al., 2015; Paganini et al., 2016; Petrou et al., 2015). On the other hand, reviews of conservation policies often fail to provide detailed information on specific SRS requirements. Only a few reviews (O’Connor et al., 2015; Petrou et al., 2015; Secades et al., 2014; Vihervaara et al., 2017) have bridged the gap between policy, remote sensing and biodiversity science. However, the current relevance of these reviews for the Kunming-Montreal GBF is limited considering their focus on national targets (Vihervaara et al., 2017) or on the previous set of Aichi targets (O’Connor et al., 2015). No detailed user requirement analysis of SRS products has yet been performed in the GBF context, which limits the capacity of the scientific community to contribute to the development of new SRS products and workflows that can be used in biodiversity indicators.

Here, we review those biodiversity indicators of the Kunming-Montreal GBF (CBD, 2022a, 2022b) that capture the state of terrestrial biodiversity, and analyse their workflows and underlying spatial information products to provide our view on key directions for development of ecologically relevant SRS products in the next decade. We use the concept of Essential Biodiversity Variables (EBVs) (Pereira et al., 2013) as an abstraction layer to sort spatial information products from the GBF indicators into major dimensions of biodiversity (i.e. variables related to species populations, species traits, community composition, ecosystem structure and ecosystem functioning). This allows us to count and score the EBV usage across GBF goals and targets, and to assess the relevance of SRS products for monitoring terrestrial biodiversity in a global policy context. Together with scores of the feasibility, accuracy and (im)maturity of SRS products from a comprehensive expert assessment (Skidmore et al., 2021), we rank SRS products and associated EBVs to create a top priority list. From this list, we provide our *viewpoint* on how the scientific community could advance biodiversity monitoring in the next decade with SRS in the context of the GBF. For clarity on the terminology, we provide a glossary of key terms used in this manuscript (see Table 1).

**Table 1.** Glossary of key terms. The list captures key terms, descriptions, and an example focused on target 2 in goal A of the Kunming-Montreal global biodiversity framework (GBF).

| Key term | Description | Example |
| --- | --- | --- |
| Goal | Four long-term goals (A, B, C, D) are defined by the Kunming-Montreal GBF (CBD, 2022a) to be achieved by 2050. | Goal A: The integrity, connectivity and resilience of all ecosystems are maintained, enhanced, or restored, substantially increasing the area of natural ecosystems by 2050. Human induced extinction of known threatened species is halted, and, by 2050, extinction rate and risk of all species are reduced tenfold, and the abundance of native wild species is increased to healthy and resilient levels. The genetic diversity within populations of wild and domesticated species, is maintained, safeguarding their adaptive potential. |
| Target | For each goal, several action-oriented conservation targets (23 in total) are defined to be met in 2030. | Target 2: Ensure that by 2030 at least 30 per cent of areas of degraded terrestrial, inland water, and coastal and marine ecosystems are under effective restoration, in order to enhance biodiversity and ecosystem functions and services, ecological integrity and connectivity. |
| Target component | In the zero and first draft of the GBF, each target is divided into target components to aid in tracking the progress towards that target. | Target 2 is divided into two target components, including T2.2 'Connectivity'. |
| Monitoring element | In the zero draft of the GBF, each target component is divided into elements to aid the selection of biodiversity indicators. | For the 'Connectivity' target component, six monitoring elements are defined, including the 'Trends in extent and rate of change of terrestrial ecosystems'. |
| Biodiversity indicator | To track the progress of each GBF goal and target, several (headline, component and complementary) biodiversity indicators are defined (CBD, 2022b). | For goal A, 22 biodiversity indicators (out of 45) are promoted by the Biodiversity Indicators Partnership (BIP) to track the state of terrestrial biodiversity, including the 'Biodiversity Habitat Index' (Hoskins et al., 2016) and the 'Species Protection Index' (GEO BON, 2015). |
| Biodiversity indicator workflow | Analytical steps (e.g., through computer-algorithms, data-processing, GIS-analyses) that are needed to process available data products (from remote sensing or in-situ observations) into biodiversity indicators. | The 'Biodiversity Habitat Index' is calculated by the 'Biogeographic modelling Infrastructure for Large-scale Biodiversity Indicators' (Hoskins et al., 2020) to map spatial patterns of the distribution of biodiversity. |
| Satellite remote sensing product | Satellite remote sensing (SRS) products are derived from satellite imagery, here referring to those which capture aspects of biodiversity and ecosystems (Skidmore et al., 2021). | The Global Forest Cover product (Hansen et al., 2013) is used (with additional biodiversity data) to calculate the 'Biodiversity Habitat Index'. |
| Essential biodiversity variable | Essential Biodiversity Variables (EBVs) are a minimum set of complementary measurements (https://geobon.org/ebvs/what-are-ebvs/; accessed on 30 January 2023), conceptually located between low-level primary observations and high-level biodiversity indicators, to capture major dimensions of biodiversity change (Pereira et al., 2013). | Examples of EBV data products are available from the GEO BON EBV data portal (https://portal.geobon.org). An example is the 'Forest loss year (2000-2018)' product as an ecosystem distribution EBV (in the EBV class 'Ecosystem structure'). |

## 2 Materials and methods

Our methodology for analysing scientific opportunities for developing SRS products in the next decade included a review of the GBF to classify and filter goals and targets that contain biodiversity indicators for monitoring the state of terrestrial biodiversity with SRS (Figure 1, top). We subsequently analysed workflows of these biodiversity indicators and summarized which spatial information products are used (Figure 1, right). By classifying each product into the EBV framework and counting the usage of different EBVs (Figure 1, bottom), we could calculate the relevance of such products in the GBF context. Together with scorings of the feasibility, accuracy and (im)maturity of SRS products by current instruments and technologies from an expert assessment (Skidmore et al., 2021), we then ranked associated EBVs on their summary scores and created a top priority list of SRS products for scientific development (Figure 1, left).

**Figure 1.**
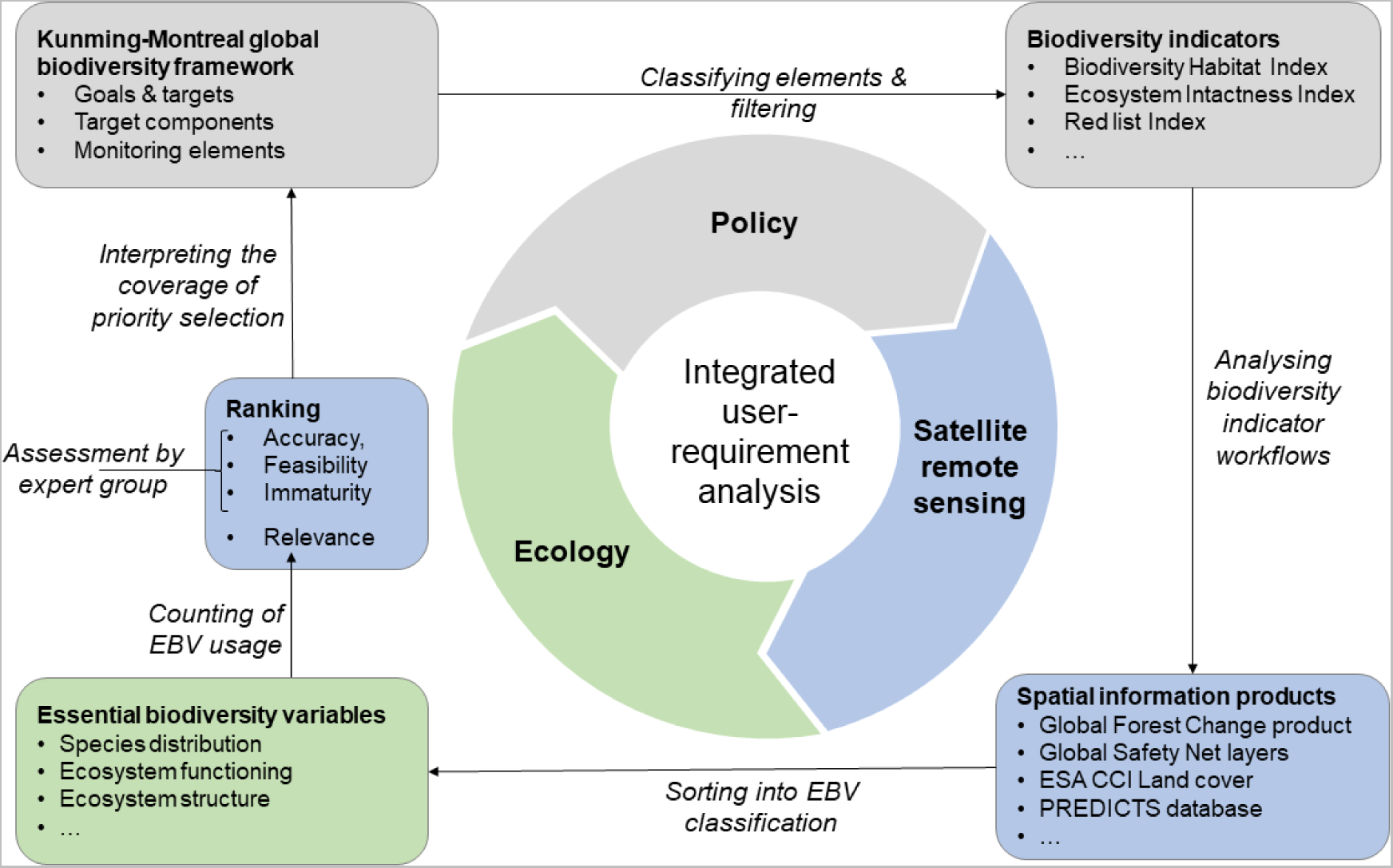
Conceptual framework of an integrated user-requirement analysis which combines policy (goals and targets of the Kunming-Montreal global biodiversity framework), satellite remote sensing (spatial information products) and ecology (essential biodiversity variables). See method section for further details.

### 2.1 Filtering of Kunming-Montreal goals and targets

The GBF defines four goals for 2050 (A–D) and 23 targets for 2030 (CBD, 2022a). A monitoring framework has been developed to track progress towards these goals and targets, composed of three groups of biodiversity indicators (i.e. headline, component and complementary) (CBD, 2022b). We used a filtering approach (Figure 1, top) to include only biodiversity indicators that capture the state of terrestrial biodiversity and which are potentially observable from space (see Table S1.2 in Supplementary Material S1). For this, we classified keywords within the descriptions of the goals and targets of the GBF (Table 1). We only included those goals and targets that capture the state of biodiversity following the Driver-Pressure-State-Impact-Response (DPSIR) framework (Burkhard and Müller, 2008).

### 2.2 Biodiversity indicator workflow analysis

For the identified goals and targets, we performed an in-depth analysis of the associated biodiversity indicators (listed in Table S3.1 of Supplementary Material S3). For each indicator, we screened the workflow as described in scientific publications, technical reports, and websites (sources in Table S3.2 of Supplementary Material S3) in terms of its relevance for tracking the state of terrestrial biodiversity. We identified all data products used in each workflow and selected those with biological geospatial information, e.g. products of biodiversity hotspots (Dinerstein et al., 2019). To provide relevant information for future SRS products, we also included spatial information products that are currently collected with methods other than SRS. Furthermore, we did not weigh SRS products by their relative importance for a given indicator workflow (e.g. whether they are required, preferred or merely desired) or by their direct or indirect use, because there is insufficient information to conduct sensitivity analyses of multiple scenarios for every workflow (Rowland et al., 2021).

Spatial information products were then sorted into the EBV classification (Figure 1, bottom). The EBV concept was useful here because it provides an intermediate (ecological) abstraction layer for connecting the spatial information products from the GBF indicators with products obtained from SRS (Figure 1). Currently, a total of 21 EBVs across six EBV classes are recognised by the Group on Earth Observations Biodiversity Observation Network (GEO BON) (https://geobon.org/ebvs/what-are-ebvs/, accessed 30 January 2023). We sorted each spatial information product into one of the 21 EBVs based on the elements of biodiversity they capture. For instance, the ESA Landcover CCI data product (ESA, 2017) was grouped into the live cover fraction EBV because it provides 38 layers of fractional cover for various ecosystem types (e.g. grassland, deciduous shrubland, lichens and mosses). Similarly, the ‘Terrestrial Ecoregions of the World’ (Sayre et al., 2020) was grouped into the ecosystem distribution EBV, as it provides information on distinct ecological assemblages (but not on fractional cover). This grouping provided us with information on how often EBVs and EBV classes are represented in the different biodiversity indicators of the GBF (full details provided in Supplementary Material S4).

### 2.3 Ranking EBV usage

The next step was to integrate the results from the workflow analysis into a ranking of specific products from current SRS instruments and technologies (Figure 1, left). For this, we first counted how often EBVs were used for each goal and target. We then applied a natural breaks classification to assign scores to the usage (1 for a usage >15, 2 for a usage >1 AND <=15, and 3 for usages ≤1). This allowed us to be consistent with the three-point scoring system used by Skidmore et al. (2021) for the other criteria. To test how our results might change if additional indicators are included that are currently insufficiently described (listed in see Table S3.3 in Supplementary Materials S3), we also performed a sensitivity analysis on this relevance score using descriptions of these (immature) indicators to specify the potential usage of EBVs (see Table S3.4 in Supplementary Materials S3).

We combined the scores for the relevance, feasibility, accuracy, and immaturity criteria (Table 2) into a summary score and created a ranking list from the summary scores to identify which SRS products could be developed by the scientific community to expand the tracking of GBF goals and targets. We therefore adapted the results from Skidmore et al. (2021) in the following way (see overview in Table 2). First, we used our relevance score (see above) to explicitly capture the relevance for the Kunming-Montreal GBF. Second, we defined immaturity as the inverse of the maturity score (Skidmore et al., 2021) to highlight the potential of (current and upcoming) SRS technologies rather than those that are currently already operational, as was the aim of Skidmore et al (2021). By combining the scores, we identified which SRS products from current instruments and technologies are most promising for future development. In the summary scores, we did not weigh the criteria to keep them equally important as well as comparable with the results of Skidmore et al. (2021). From the ranking of the summary scores, we then identified the top priority products by focusing on the upper 10^th^ percentile of the list, corresponding to a threshold of a summary score equal to 7.

**Table 2.** Criteria used for scoring satellite remote sensing products in the context of the Kunming-Montreal global biodiversity framework (GBF). See method section for further details.

| Criteria | Description | Score (good = 1) | Score (poor = 3) |
| --- | --- | --- | --- |
| Relevance | A measure of how satellite remote sensing products with a specific biodiversity focus (i.e. related to Essential Biodiversity Variables, EBVs) could contribute to the GBF 2030 targets and goals. | $= \begin{cases} 1 & \text{for } EBV_{usage} > 15 \\ 2 & \text{for } EBV_{usage} > 1 \text{ \& } EBV_{usage} \leq 15 \\ 3 & \text{for } EBV_{usage} \leq 1 \end{cases}$ | |
| Feasibility | The science community knows how to measure the remote sensing biodiversity product at appropriate scales that such measurements can realistically be made and/or that observations already exist | Indicates the maturity of science/ technology/ experience needed to make the remote sensing biodiversity product | Indicates that considerable research and development effort remain or that remote sensing biodiversity products at the scale needed are technically, logistically or financially difficult to make |
| Accuracy | A measure of the accurate observation of a given remote sensing variable. It considers the effectiveness of remote sensing data and techniques to achieve an accurate and precise value of the remote sensing enabled biodiversity variable. | Fully operational network or service is in place, generating remote sensing biodiversity products that are accurate for the purpose | Indicates that no or very limited action has been taken to generate accurate remote sensing biodiversity products |
| Immaturity | The potential of developing remote sensing biodiversity products can be identified and/or proposed to a funding body. | Indicates that at present a relevant infrastructure for the remote sensing variable is absent and consequently still has a high potential to be operationally produced. | Indicates the remote sensing variable is mature (with little development steps required) and is operationally produced within a relevant infrastructure. |

## 3 Results

### 3.1 Filtering of Kunming-Montreal goals and targets

Across the goals and targets of the Kunming-Montreal GBF (CBD, 2022a), we identified 54 keywords (see Table S1.1 in Supplementary Materials S1). Following the classification of these keywords, we identified 5 keywords (see Table S1.2 in Supplementary Materials S1) to be drivers of biodiversity (e.g., agriculture, harvest), 8 to be pressures (e.g., acidification, extinction), 11 to represent the state (e.g., integrity, connectivity and resilience), 8 to characterise the impact (e.g., contributions to human, utilization), and 22 as responses to biodiversity loss (e.g., adaptation, implementation, policy). By selecting only those goals and targets that pertain to the state of the biodiversity, we identified two GBF goals and eight targets (see Table 3) of which targets T01, T02, T03, T06 and T08 aimed at goal A and T10, T11 and T12 at goal B. This selection was consistent with a previous analysis performed by us on the monitoring elements of the updated zero draft version of the GBF (see Figure S1.1, Table S1.2, and Table S1.3 of Supplementary Materials S1). None of the targets in goals C and D focused on monitoring the state of terrestrial biodiversity with SRS observations, as these goals focus on genetic resources (goal C) and management practises (goal D). For the identified goals and targets, we found 87 usages of (non-unique) terrestrial and agnostic biodiversity indicators, spread over goal A (*n* = 32) and goal B (*n* = 8) and specific targets (*n* = 47) (see Table S3.1 in Supplementary Material S3). We also found that some indicators are used multiple times within different goals and targets, such as the Bioclimatic Ecosystem Resilience Index (2×) and the Red list Index (9×), finally leading to a total of 58 unique indicators.

**Table 3.** Summary descriptions of the goals and target components of the Kunming-Montreal global biodiversity framework (GBF) which aim to monitor the state of terrestrial biodiversity. The eight targets (T) are spread over two goals (A, B) and were identified by analysing the full descriptions of the GBF targets and goals, provided in Table S1.1 of Supplementary Material S1.

| Goal description | Target description |
| --- | --- |
| Goal A: Reduce threats to habitat and wild and domesticated species. | T01. Reduce the loss of areas of high biodiversity importance to zero. |
|  | T02. Improve connectivity of priority terrestrial ecosystems. |
|  | T03. Ensure management of protected areas |
|  | T06. Control introduction and impacts of invasive alien species. |
|  | T08. Minimize the impact of climate change. |
| Goal B: Meet people's needs by improving the sustainable use of ecosystem functions and services | T10. Increase sustainable productivity of agriculture and forested ecosystems. |
|  | T11. Enhance nature's contributions to people. |
|  | T12. Increase public urban green/blue spaces |

### 3.2 Biodiversity indicator workflow analysis

Of the 58 unique biodiversity indicators, 36 provided sufficient information on their workflows (details in Table S3.2 in Supplementary Material S3), whereas 22 were insufficiently described because they are still in development (see Table S3.3 in Supplementary Material S3). From the 36 workflows providing sufficient information, we identified 95 spatial information products (details in Table S4.1 in Supplementary Material S4).

This included, for instance, the Global Forest Change product (Hansen et al., 2013) and the CCI landcover product (ESA, 2017). Each spatial information product was then classified into one of the 21 EBVs (see detailed overview in Supplementary Material S4). Products associated with the EBV classes ‘species populations’ and ‘ecosystem structure’ were most often used (Figure 3). Within the ‘ecosystem structure’ EBV class, the highest usage was related to the ecosystem distribution EBV and live cover fraction EBV (Figure 3). Within the ‘species populations’ EBV class, most spatial information products were related to the species distributions EBV and less to the species abundances EBV (Figure 3).

**Figure 3:**
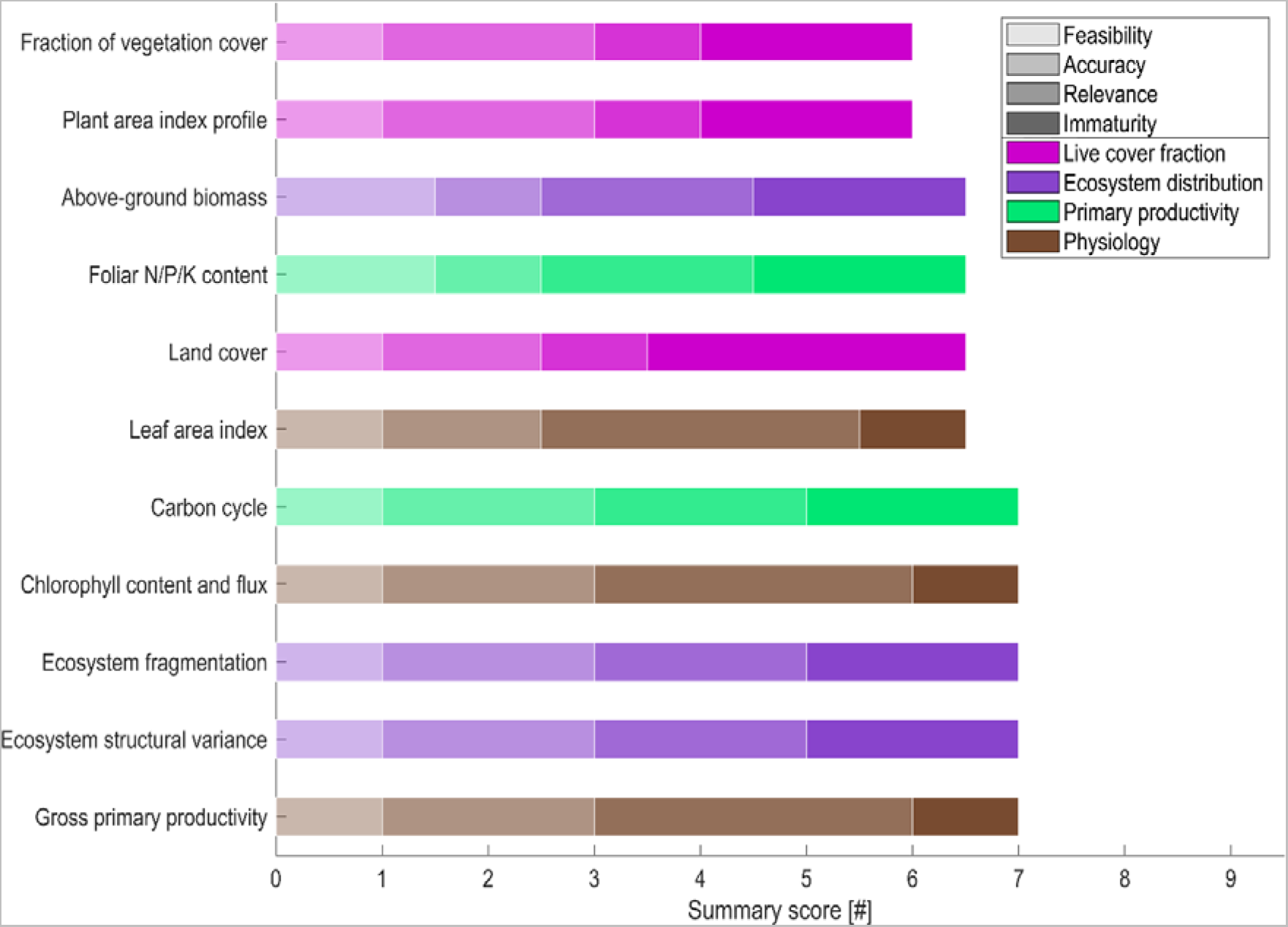
Priority list showing the satellite remote sensing products with the best summary score and the highest rank. The best ranking reflects those products that have the lowest summary score. The colours of the bars denote the grouping into the Essential Biodiversity Variable framework, and the brightness of the bars indicate the scores of the feasibility, accuracy, immaturity and relevance criteria (ranging from 1 to 3). See Table S5.1 in Supplementary Material S5 for the scores and ranking of all 56 satellite remote sensing products.

### 3.3 Ranking EBV usage

Products of live cover fraction, ecosystem distribution, and species distributions EBVs were most often used in the GBF (Figure 2), which resulted in a high relevance score (= 1). In contrast, products associated with the EBV classes ‘community composition’ (e.g. community abundance) and ‘ecosystem functioning’ (e.g. primary productivity and ecosystem disturbance) were moderately used by the identified biodiversity indicators and therefore given a moderate relevance score (= 2). All other EBVs received a low relevance score (= 3) (see numbers at bottom of Figure 2).

**Figure 2.**
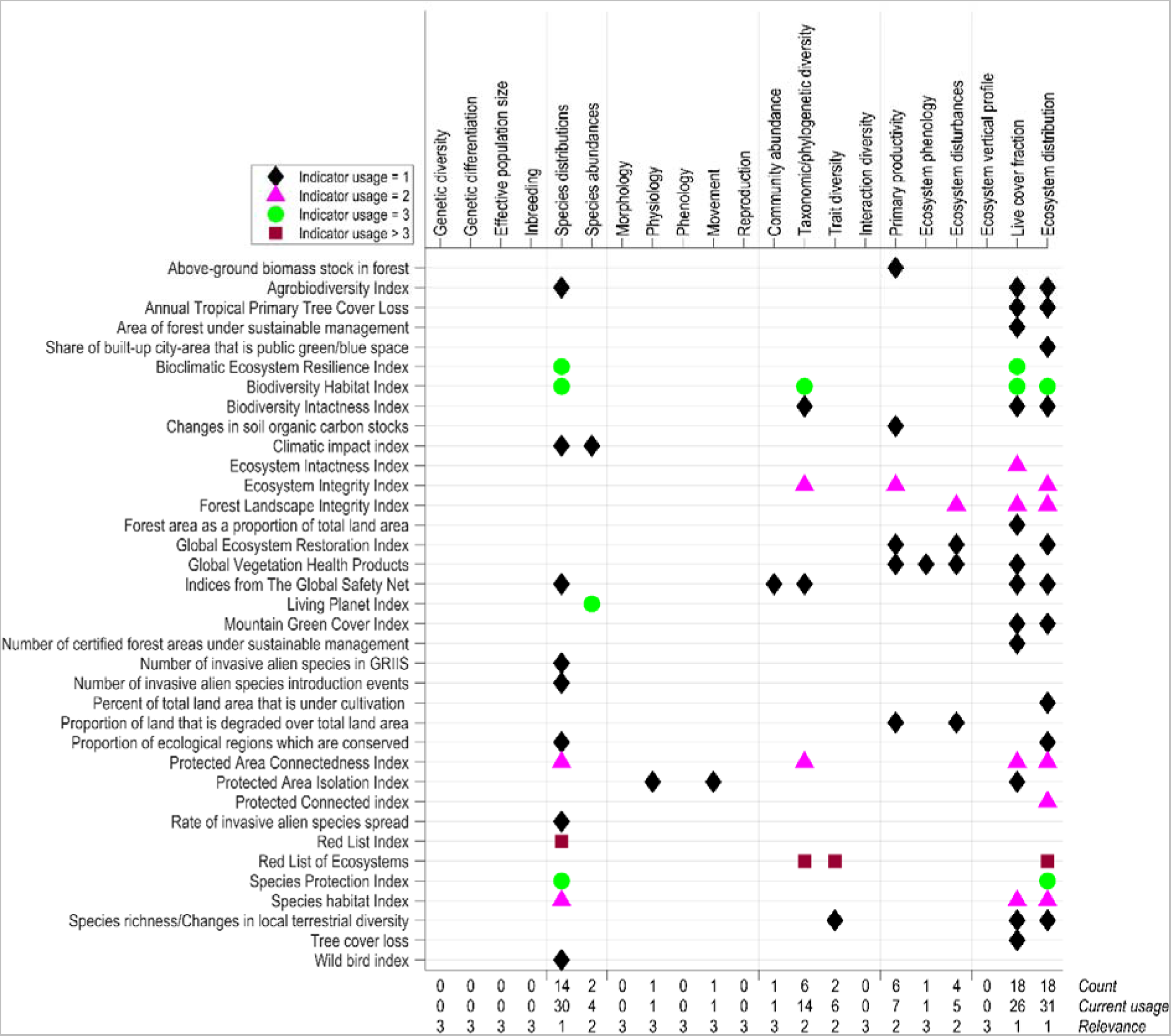
Biodiversity indicators and their underlying data categorised into Essential Biodiversity Variables (EBVs). Full names of the biodiversity indicators are provided in Table S3.2 in Supplementary Material S3. The indicators (y-axis) represent those listed in the Kunming-Montreal global biodiversity framework (GBF) for goals and targets that focus on tracking the state of terrestrial biodiversity. As some indicators are used several times by different goals and targets (e.g. Red List Index 9×, Red list of Ecosystems 5×), we use the following symbol colours: black diamonds = 1, purple triangles = 2, green circles = 3, brown squares > 3. At the bottom, we calculate how often EBVs are used by the different biodiversity indicators individually (count), and how often across the selected GBF targets and goals (current usage). This current usage is then scored (see Table 2) to describe the relevance of these EBVs across the GBF (relevance).

The sensitivity analysis showed that relevance scores remained comparable when including the 22 insufficiently documented indicators (see Figure S3.1 in Supplementary Material S3). Only data products associated with the ecosystem phenology EBV improved their relevance score from 3 (poor) to 2 (moderate), but this did not have major impacts on the ranking and prioritization of the SRS products (Table S5.1 in Supplementary Material S5).

Eleven SRS products were ranked highest after combining the relevance scores with the feasibility, accuracy, and immaturity scores into a summary score (Figure 3). The SRS products with the highest rank (summary score = 6) were fraction of vegetation cover and plant area index profile, followed by above-ground biomass, foliar N/P/K content, land cover (vegetation type) and leaf area index (summary score = 6.5). Five additional SRS products were also included in the top priority list (summary score = 7), namely carbon cycle biomass (above-ground biomass), chlorophyll content and flux, ecosystem fragmentation, ecosystem structural variance, and gross primary productivity (Figure 2).

## 4 Discussion

Satellite remote sensing plays a vital role for monitoring the pressures, impacts and drivers of biodiversity change (Lehmann et al., 2022; Masó et al., 2020), but the role of SRS for directly monitoring the state of biodiversity remains less clear. Here, we reviewed the goals, targets and biodiversity indicators of the Kunming-Montreal GBF to provide our viewpoint on how SRS products could be improved for tracking the state of terrestrial biodiversity from space. We identified a priority list of SRS products for further development by the scientific community which could enhance the monitoring of GBF goals and targets. Our priority list contains SRS products that would benefit from improvements for their use in the GBF (e.g. fractional cover, landcover, above ground biomass and gross primary production), but also immature SRS products that are not yet used (e.g. foliar N/P/K content) and products for which alternatives and proxies are currently in place (e.g. NDVI for vegetation coverage by leaf area index). From our analysis, we suggest two key directions for advancing SRS products and workflows in the context of the GBF, namely (1) the refinement of SRS products (derived from current instruments) associated with ecosystem distribution, live cover fraction and species distributions, and (2) the creation of new SRS products (from current and upcoming technologies) associated with plant physiology and primary productivity.

Our review reveals that half of the analysed GBF indicators (i.e. 18 out of 36) require geospatial information about ecosystem distributions, such as the Global Forest Change product (Hansen et al., 2013). The data sources currently used in the GBF indicator workflows are diverse, with strong differences in underlying methodology and spatial resolution, e.g. in radiometric correction, classification, legend information and grain size (Herold et al., 2008). This can potentially lead to inconsistencies between indicators (Martin et al., 2019; Tuanmu and Jetz, 2014) and consequently affect their comparability and reliability for tracking the GBF goals and targets. We therefore suggest that ecosystem distribution products need to be improved to become more consistent, more harmonized, and more ecologically relevant. For instance, land use information often remains too coarse (e.g. 1 km resolution) for ecological applications, as many biodiversity indicators —such as the Species Protection Index (GEO BON, 2015) or the Bioclimatic Ecosystem Resilience Index (Ferrier et al., 2020)— require finer spatial resolutions (e.g. 10–30 m). While high resolution SRS products exist, e.g. GlobLand30 (Chen et al., 2015), Google’s Dynamic World (Brown et al., 2022), or ESA’s World Cover (Zanaga et al., 2021), their thematic detail and typology are insufficient for tracking changes in ecosystem distributions. Hence, a large scientific opportunity exists for refining and improving such products, e.g. by developing globally consistent typologies for ecosystem distribution maps (Keith et al., 2022). Moreover, combining SRS products of land cover, vegetation type or function, soil types, precipitation and elevation with in-situ observations of plant species (e.g. from vegetation plots or citizen science collections) will allow to create suitable ecosystem distribution maps, especially through applications of deep-learning models that include high resolution SRS products with high temporal frequency to capture the spatial structure of landscapes (Estopinan et al., 2022).

About half of the biodiversity indicators of the GBF also use geospatial data products that are associated with the live cover fraction EBV. Live cover fraction products derived from SRS, such as vegetation fraction from MODIS (Dimiceli et al., 2015), are already used in biodiversity indicators to characterize ecosystem quality (Venter et al., 2016), ecosystem integrity (Grantham et al., 2020) and habitat intactness (GEO BON, 2015). The potential of SRS to estimate live cover fraction has also been widely recognised (Gao et al., 2020; Skidmore et al., 2015; Valbuena et al., 2020; Vihervaara et al., 2017), but current SRS products do not allow to relate the fraction of each vegetation (or plant functional) type to an ecosystem reference state (e.g. based on an ecosystem taxonomy). Most SRS products of fractional cover, such as the ESA CCI product (ESA, 2017), have been developed for climate change modelling and hence are not detailed enough for defining ecosystems in the context of biodiversity indicators. Hence, it remains understudied how the state of fractional cover (e.g. of heath, shrubs, grasses or other growth forms) relates to different ecosystem taxonomies as used in biodiversity indicators, such as the terrestrial ecoregions of the world (Olson et al., 2001) or the terrestrial habitat classification of the European Union (Chytrý et al., 2020). Development of better ecosystem live cover fraction products will therefore benefit from close collaboration between SRS scientists, ecologists and policy experts (Pettorelli et al., 2016). We encourage the co-design and co-production of such products together with large-scale biodiversity research e-infrastructures (e.g. GBIF, LifeWatch ERIC), space agencies (e.g. NASA and ESA), biodiversity observation networks (e.g. GEO BON), and parties developing National Biodiversity Strategies and Action Plans (NBSAPs) to ensure an effective implementation of the GBF goals and targets (Xu et al., 2021).

More than one third of the analysed biodiversity indicators of the GBF (i.e. 40%) use spatial information products associated with the species distributions EBV, e.g. the Living Planet Index (Collen et al., 2013) or the Red List (Butchart et al., 2007). These products are often needed in the context of monitoring threatened, range-restricted species, alien invasive species or areas of conservation importance. However, SRS is rarely used for directly mapping species distributions because spatial resolutions of openly accessible products are most often not high enough for individual species detection (Sladonja and Damijanić, 2021). While classifications of native and invasive plant species in tropical forests (Asner et al., 2017; Somers and Asner, 2013) have been performed with imaging spectroscopy data, these applications usually use airborne (rather than spaceborne) observations. Using SRS for directly mapping species distributions is thus unlikely to become operational in the near future unless spaceborne instruments with very high resolutions will become publicly and freely available. Hence, the most obvious opportunities for using SRS observations for mapping species distributions are related to providing co-variates in species distributions models (Randin et al., 2020), rather than detecting species directly from space. In this context, we see the most promising advances in SRS through sensors and products that can better capture the environmental condition of animal habitats (Gudex-Cross et al., 2021), the environmental conditions (Radeloff et al., 2019), and structural heterogeneity derived from a combination of LiDAR, radar and multispectral observations (Valbuena et al., 2020).

Current biodiversity indicators often omit SRS products related to plant physiology (e.g. leaf area index and chlorophyll content) and primary productivity (e.g. foliar N/P/K). However, such SRS products could provide key information on biodiversity change, e.g. on plant phenology (e.g. timing of flowering and fruiting) and stoichiometry (e.g. the balance between carbon, nitrogen and phosphorus) (Kissling et al., 2018b), grassland carbon sequestration (Martini et al., 2019), ecosystem resilience (Longo et al., 2018; Schneider et al., 2023) or forest net primary productivity (Šímová et al., 2019). Information on (canopy and leaf) trait dissimilarities further allows to discriminate alien invasive species (Große-Stoltenberg et al., 2016), their impact on ecosystem functioning (Große-Stoltenberg et al., 2018), nutrient cycling (Kumar and Garkoti, 2021) and energy exchange (Musavi et al., 2015). We therefore advocate the creation of scalable workflows to produce new SRS products that can characterise individual-level plant functioning (via physiology), which subsequently can be aggregated at the ecosystem level (e.g. primary productivity). At present, most satellite observations remain too coarse (≥ 20 m) in their spatial resolution for detecting individual species (Kissling et al., 2018a; Skidmore et al., 2021). While satellites with higher spatial resolutions (< 2 m) exist, e.g. Superview and Worldview, they have limited temporal and/or spectral capabilities that currently prohibit the estimation of physiological plant traits at the species level. One promising opportunity would be to merge multiple satellite observations for estimating primary productivity and plant physiology at the plant level (Reddy et al., 2021), not just for leaf area index, chlorophyll content and foliar nitrogen, but also for other plant functional traits (e.g., leaf mass area and water content). Deep-learning techniques such as DeepForest can already be applied to map individual trees (Brandt et al., 2020; Weinstein et al., 2021), while 3D models such as DART can be used to characterise plant heterogeneity within coarser pixels (Gastellu-Etchegorry et al., 2004). Similarly, land surface models such as BETHY-SCOPE can be used to simulate the gross uptake of carbon from optical, fluorescent and thermal observations (Norton et al., 2019). Using multi-sensor frameworks (Lewis et al., 2012) and data science techniques, such as model emulators (Gómez-Dans et al., 2016; Verrelst et al., 2012) and neural networks (Baret et al., 2013; Cherif et al., 2023), would allow to integrate the above-mentioned components into consistent and scalable products of plant traits and primary productivity. This would allow the estimation of leaf area index, foliar chlorophyll and nitrogen content with hyperspectral observations (Berger et al., 2020; Féret et al., 2021) and even with multi-spectral observations (Bossung et al., 2022), which could allow to better track plant species phenology, vegetation health, alien invasive species, and the consequences of biodiversity change on ecosystem functioning.

Capitalizing on the abovementioned opportunities for developing SRS products could make important contributions to the Kunming-Montreal GBF in the next decade and the subsequent period of 2030–2050. Until 2030, the biggest opportunities are those that employ new SRS products from current instruments and technologies in indicator workflows to track goal A and B, and targets T1, T2, T3, T8, T10, T11 and T12 (Figure 4). For instance, refining ecosystem distribution and live cover fraction products could greatly enhance the tracking of goal A (‘reducing species extinction rates’) as well as target 2 (‘improve connectivity of priority terrestrial ecosystems) and target 3 (‘ensure management of protected areas’), as these targets already involve biodiversity indicators which use such products (see Figure 4). For SRS products related to primary productivity and plant species physiology, novel products could innovate biodiversity indicators to allow better tracking of alien invasive species in target 6 (‘control the introduction and impact of invasive alien species’), characterise ecosystem resistance to climate change in target 8 (‘minimize impact of climate change on biodiversity’) as well as a better estimation of primary productivity from plant functional traits for target 10 (‘increase sustainable productivity of agricultural and forested ecosystems’). Such new SRS products of plant physiology and primary productivity and improved ecosystem distribution and live cover fraction products (Schimel et al., 2019) could also be used as predictors for modelling species distributions which are broadly needed across most of the GBF (see Figure S4.1 in Supplementary Materials S4).

**Figure 4.**
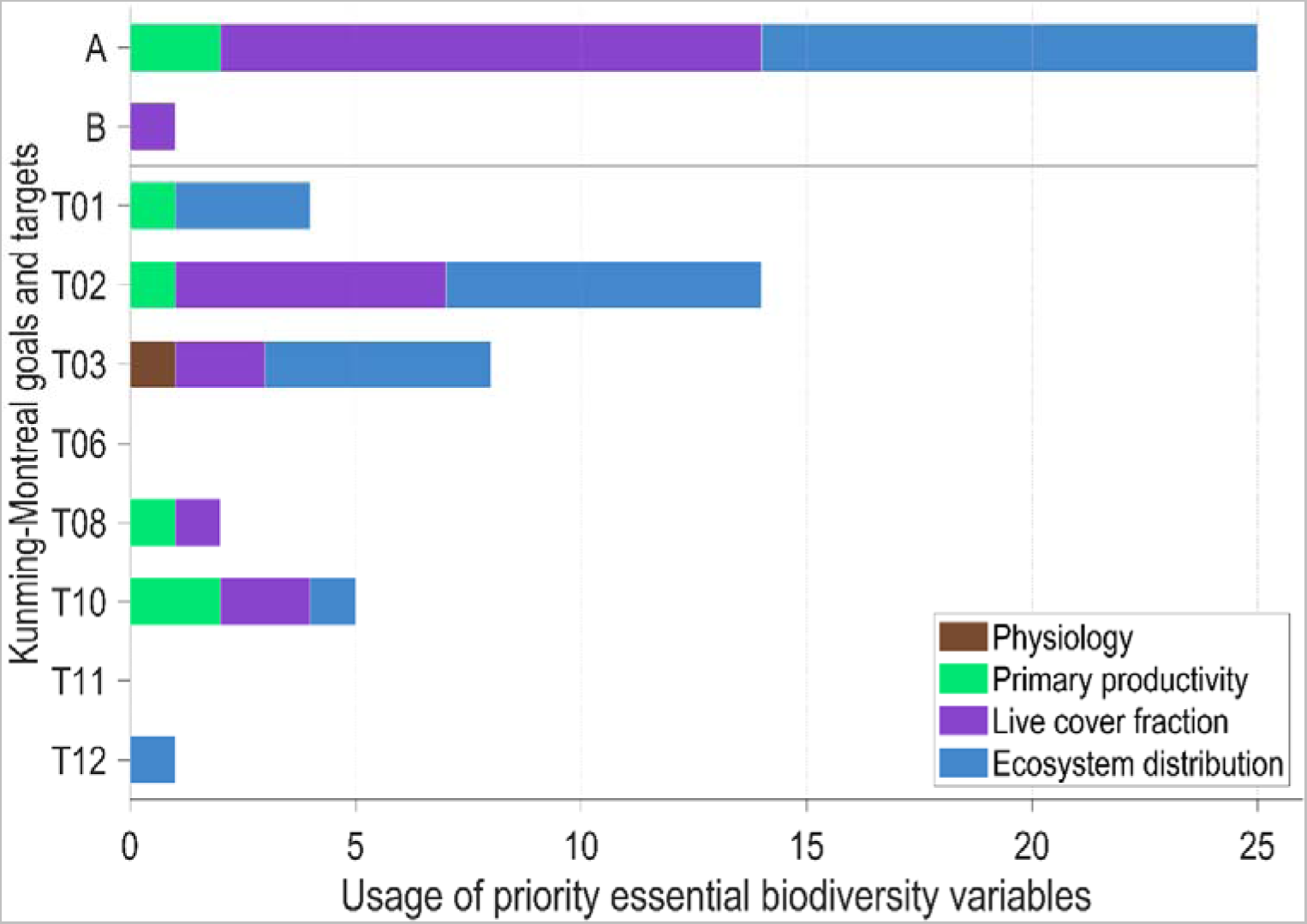
The usage of priority essential biodiversity variables (EBVs) derived from space in the context of the goals and targets of the Kunming-Montreal global biodiversity framework. See Table 3 for abbreviations and summary descriptions of the goals (A, B) and targets (T).

For the period 2030–2050, additional improvements in tracking the goals for 2050 can be accomplished if biodiversity indicators utilize the capabilities of new satellite instruments and missions. With advances in the capabilities of new satellites —such as higher spatial resolutions (Sadeh et al., 2021), new spectral capabilities (Coppo et al., 2017) and laser scanning (Dubayah et al., 2020)— SRS can satisfy user needs that are currently not met by existing technologies and products. For example, developments in tracking animal movement and population sizes from space can provide new potential for SRS products and biodiversity indicators to be developed (Jetz et al., 2022; Lahoz-Monfort and Magrath, 2021; Wu et al., 2023). While the movement EBV and other EBVs (e.g., trait diversity) currently have a low relevance in the GBF (see Figure 2), this is mainly driven by data being not available for implementation into biodiversity indicators. To define scientific opportunities for the period after 2030, our approach needs to be extended to identify mission requirements, for example by adapting the approach used in NASA’s Decadal Survey to link driving scientific questions to measurement requirements and geophysical observables for their Surface Biology and Geology (SBG) observatory (Schimel et al., 2020; Stavros et al., 2023). By including a broad community input (e.g., from the CBD’s Ad Hoc Technical Expert Group on Indicators), such an approach can ensure to identify needs, gaps and requirements of (additional) biodiversity indicators (in development) that are broadly supported and agreed on. By advancing SRS products of current instruments and technologies for monitoring the 2030 targets and simultaneously guiding new technologies for the 2050 goals, SRS can enable a transformative and lasting change in tracking and monitoring biodiversity and ecosystems at a global scale.

## 5 Conclusion

In our *viewpoint* article, we have reviewed the biodiversity indicators of the Kunming-Montreal GBF for tracking the state of terrestrial biodiversity. Using EBVs as a broker between biodiversity policy and remote sensing, we identified several opportunities for the scientific community to advance terrestrial biodiversity monitoring from space in the current decade. Our approach thereby demonstrates how policy goals and targets can be translated into scientific opportunities. This could be expanded beyond the state of terrestrial biodiversity, e.g. to aquatic habitats (e.g. marine, coastal and freshwater systems) or other DPSIR classes (i.e. drivers, pressures, impacts and responses), as well as to other national or international policy documents that contain conservation goals and targets. We hope that this *viewpoint* inspires more collaboration between policy experts, ecologist and remote sensing scientists to advance the tracking of policy goals and targets and to contribute to reducing the threats to biodiversity while meeting people’s needs through the sustainable use of biodiversity.

## 6 Acknowledgements

We acknowledge the critical and constructive feedback from five reviewers. We are also grateful for discussions with members of the Biogeography & Macroecology (BIOMAC) lab at the University of Amsterdam, specifically Harry Seijmonsbergen, Kenneth Rijsdijk, Stacy Shinneman, and Yifang Shi. We acknowledge support from LifeWatch (https://www.lifewatch.eu/), discussions within the Group on Earth Observations Biodiversity Observation Network (GEO BON), and stimulating exchange with the members and partners contributing to the co-design of the European Biodiversity Observation Network (EuropaBON).

## 7 CRediT authorship contribution statement

**Joris Timmermans:** Conceptualization, Methodology, Validation, Formal analysis, Writing – original draft, Writing – review & editing, Visualization.

**W. Daniel Kissling:** Conceptualization, Formal analysis, Validation, Writing – original draft, Writing – review & editing, Visualization, Supervision, Funding acquisition.

