## Supplementary Materials S1 for "Advancing terrestrial biodiversity monitoring with satellite remote sensing in the context of the Kunming-Montreal global biodiversity framework"

### Supplementary Material S1

**Table S1.1. Full description of the goals and targets of the Kunming-Montreal global biodiversity framework of the Convention of Biological Diversity (CBD, 2020). For each goal and target, the main objective is identified (with underlined text). Keywords of these objectives are classified according into the DPSIR classes (Burkhard and Müller, 2008) and accordingly coloured (orange = Driver, red = Pressure, green = State, black = Impact, purple = Response). An * identifies those goals and targets that pertain to the state of biodiversity and consequently are selected for further analysis. An overview of the keyword classification is provided in Table S1.2.**

**GOAL A***

- The **integrity**, **connectivity** and **resilience** of all ecosystems are **maintained**, **enhanced**, or **restored**, substantially increasing the **area of natural ecosystems** by 2050;
- Human induced **extinction** of known threatened species is halted, and, by 2050, **extinction** rate and **risk** of all species are reduced tenfold, and the abundance of native wild species is increased to healthy and **resilient** levels;
- The genetic diversity within populations of wild and domesticated species, is maintained, safeguarding their adaptive potential.

**GOAL B***

Biodiversity is sustainably used and managed and **nature’s contributions to people**, including **ecosystem functions** and **services**, are valued, **maintained** and **enhanced**, with those currently in decline being **restored**, supporting the achievement of sustainable development, for the benefit of present and future generations by 2050

**GOAL C**

The **monetary** and **non-monetary** benefits from the **utilization** of genetic resources, and digital sequence information on genetic resources, and of traditional knowledge associated with genetic resources, as applicable, are shared **fairly** and **equitably**, including, as appropriate with indigenous peoples and local communities, and substantially increased by 2050, while ensuring traditional knowledge associated with genetic resources is appropriately protected, thereby contributing to the conservation and sustainable use of biodiversity, in accordance with internationally agreed access and benefit-sharing instruments.

**GOAL D**

Adequate means of **implementation**, including financial resources, **capacity building**, technical and scientific **cooperation**, and access to and transfer of technology to fully implement the Kunming-Montreal global biodiversity framework are secured and **equitably** accessible to all Parties, especially developing countries, in particular the least developed countries and small island developing States, as well as countries with economies in transition, progressively closing the biodiversity finance gap of $700 billion per year, and aligning financial flows with the Kunming-Montreal Global Biodiversity Framework and the 2050 Vision for Biodiversity

**TARGET 1***

Ensure that all areas are under participatory integrated biodiversity inclusive spatial **planning** and/or effective **management** processes addressing land and sea use change, to bring the loss of **areas of** **high biodiversity importance**, including ecosystems of high **ecological integrity**, close to zero by 2030, while respecting the rights of indigenous peoples and local communities,

**TARGET 2***

Ensure that by 2030 at least 30 per cent of areas of degraded terrestrial, inland water, and coastal and marine ecosystems are under effective **restoration**, in order to enhance biodiversity and **ecosystem functions** and **services**, ecological **integrity** and **connectivity**.

**TARGET 3***

Ensure and enable that by 2030 at least 30 per cent of terrestrial, inland water, and of coastal and marine areas, especially **areas of particular importance for biodiversity** and **ecosystem functions** and **services**, are effectively **conserved** and **managed** through ecologically representative, well-connected and **equitably** governed systems of protected areas and other effective area-based conservation measures, recognizing indigenous and traditional territories, where applicable, and integrated into wider landscapes, seascapes and the ocean, while ensuring that any sustainable use, where appropriate in such areas, is fully consistent with conservation outcomes, recognizing and respecting the rights of indigenous peoples and local communities including over their traditional territories, Ensure urgent management actions, to halt human induced extinction of known threatened species and for the recovery and conservation of species, in particular threatened species, to significantly reduce extinction risk, as well as to maintain and restore the genetic diversity within and between populations of native, wild and domesticated species to maintain their adaptive potential, including through in situ and ex situ conservation and sustainable management practices, and effectively manage human-wildlife interactions to minimize human-wildlife conflict for coexistence.

**TARGET 4**

Ensure urgent **management** actions, to halt human induced **extinction** of known threatened species and for the **recovery** and **conservation** of species, in particular threatened species, to significantly reduce **extinction** **risk**, as well as to maintain and restore the genetic diversity within and between populations of native, wild and domesticated species to maintain their adaptive potential, including through in situ and ex situ conservation and sustainable management practices, and effectively manage human-wildlife interactions to minimize human-wildlife conflict for coexistence.

**TARGET 5**

Ensure that the use, **harvesting and trade of wild species** is **sustainable**, safe and **legal**, preventing overexploitation, minimizing impacts on non-target species and ecosystems, and reducing the risk of pathogen spill-over, applying the ecosystem approach, while respecting and protecting customary sustainable use by indigenous peoples and local communities.

**TARGET 6***

Eliminate, minimize, reduce and or mitigate the impacts of **invasive alien species** on **biodiversity** and **ecosystem services** by identifying and managing **pathways** of the introduction of alien species, preventing the introduction and establishment of priority **invasive alien species**, reducing the rates of introduction and establishment of other known or potential **invasive alien species** by at least 50 per cent, by 2030, **eradicating** or **controlling** invasive alien species especially in **priority sites**, such as islands.

**TARGET 7**

Reduce **pollution** **risks** and the negative impact of **pollution** from all sources, by 2030, to levels that are not harmful to biodiversity and ecosystem functions and services, considering cumulative effects, including: reducing excess nutrients lost to the environment by at least half including through more efficient nutrient cycling and use; reducing the overall risk from pesticides and highly hazardous chemicals by at least half including through integrated pest management, based on science, taking into account food security and livelihoods; and also preventing, reducing, and working towards eliminating plastic pollution.

**TARGET 8***

Minimize the impact of climate change and ocean **acidification** on biodiversity and increase its **resilience** through mitigation, adaptation, and disaster risk reduction actions, including through nature-based solution and/or ecosystem based approaches, while minimizing negative and fostering positive impacts of climate action on biodiversity.

**TARGET 9**

Ensure that the **management** and use of wild species are **sustainable**, thereby providing social, economic and environmental benefits for people, especially those in vulnerable situations and those most dependent on biodiversity, including through sustainable biodiversity-based activities, products and services that enhance biodiversity, and protecting and encouraging customary sustainable use by indigenous peoples and local communities.

**TARGET 10***

Ensure that **areas** under agriculture, aquaculture, fisheries and forestry are **managed** **sustainably**, in particular through the sustainable use of biodiversity, including through a substantial increase of the application of biodiversity friendly practices, such as sustainable intensification, agroecological and other innovative approaches contributing to the resilience and long-term efficiency and productivity of these production systems and to food security, conserving and restoring biodiversity and maintaining nature’s contributions to people, including ecosystem functions and services.

**TARGET 11***

Restore, **maintain** and **enhance** **nature’s contributions to people**, including **ecosystem functions** and **services**, such as regulation of air, water, and climate, soil health, pollination and reduction of disease risk, as well as protection from natural hazards and disasters, through nature-based solutions and ecosystem based approaches for the benefit of all people and nature.

**TARGET 12***

Significantly increase the **area** and **quality** and **connectivity** of, access to, and benefits from **green and blue spaces** **in urban and** **densely populated** areas sustainably, by mainstreaming the conservation and sustainable use of biodiversity, and ensure biodiversity-inclusive urban planning, enhancing native biodiversity, ecological connectivity and integrity, and improving human health and well-being and connection to nature and contributing to inclusive and sustainable urbanization and the provision of ecosystem functions and services.

**TARGET 13**

Take effective **legal**, **policy**, **administrative** and **capacity**-**building** **measures** at all levels, as appropriate, to ensure the fair and equitable sharing of benefits that arise from the utilization of genetic resources and from digital sequence information on genetic resources, as well as traditional knowledge associated with genetic resources, and facilitating appropriate access to genetic resources, and by 2030 facilitating a significant increase of the benefits shared, in accordance with applicable international access and benefit-sharing instruments.

**TARGET 14**

Ensure the full integration of biodiversity and its multiple values into **policies**, **regulations**, **planning** and development processes, poverty eradication **strategies**, **strategic** environmental assessments, environmental impact assessments and, as appropriate, national accounting, within and across all levels of government and across all sectors, in particular those with significant impacts on biodiversity, progressively aligning all relevant public and private activities, fiscal and financial flows with the goals and targets of this framework.

**TARGET 15**

Take **legal**, **administrative** or **policy** **measures** to encourage and enable business, and in particular to ensure that large and transnational companies and financial institutions:

(a) Regularly monitor, assess, and transparently disclose their risks, dependencies and impacts on biodiversity including with requirements for all large as well as transnational companies and financial institutions along their operations, supply and value chains and portfolios;

(b) Provide information needed to consumers to promote sustainable consumption patterns;

(c) Report on compliance with access and benefit-sharing regulations and measures, as applicable;

in order to progressively reduce negative impacts on biodiversity, increase positive impacts, reduce biodiversity-related risks to business and financial institutions, and promote actions to ensure sustainable patterns of production.

**TARGET 16**

Ensure that people are encouraged and enabled to make **sustainable** **consumption** choices including by establishing supportive policy, legislative or regulatory frameworks, improving education and access to relevant and accurate information and alternatives, and by 2030, reduce the global footprint of consumption in an equitable manner, halve global food waste, significantly reduce overconsumption and substantially reduce waste generation, in order for all people to live well in harmony with Mother Earth.

**TARGET 17**

Establish, **strengthen** **capacity** for, and implement in all countries in biosafety **measures** as set out in Article 8(g) of the Convention on Biological Diversity and measures for the handling of biotechnology and distribution of its benefits as set out in Article 19 of the Convention.

**TARGET 18**

Identify by 2025, and **eliminate**, phase out or reform incentives, including subsidies harmful for biodiversity, in a proportionate, just, fair, effective and equitable way, while substantially and progressively reducing them by at least 500 billion United States dollars per year by 2030, starting with the most harmful incentives, and scale up positive incentives for the conservation and sustainable use of biodiversity.

**TARGET 19**

Substantially and progressively increase the level of **financial** resources from all sources, in an effective, timely and easily accessible manner, including domestic, international, public and private resources, in accordance with Article 20 of the Convention, to implement national biodiversity strategies and action plans, by 2030 mobilizing at least 200 billion United States dollars per year, including by:

(a) Increasing total biodiversity related international financial resources from developed countries, including official development assistance, and from countries that voluntarily assume obligations of developed country Parties, to developing countries, in particular the least developed countries and small island developing States, as well as countries with economies in transition, to at least US$ 20 billion per year by 2025, and to at least US$ 30 billion per year by 2030;

(b) Significantly increasing domestic resource mobilization, facilitated by the preparation and implementation of national biodiversity finance plans or similar instruments according to national needs, priorities and circumstances

(c) Leveraging private finance, promoting blended finance, implementing strategies for raising new and additional resources, and encouraging the private sector to invest in biodiversity, including through impact funds and other instruments;

(d) Stimulating innovative schemes such as payment for ecosystem services, green bonds, biodiversity offsets and credits, benefit-sharing mechanisms, with environmental and social safeguards

(e) Optimizing co-benefits and synergies of finance targeting the biodiversity and climate crises,

(f) Enhancing the role of collective actions, including by indigenous peoples and local communities, Mother Earth centric actions and non-market-based approaches including community based natural resource management and civil society cooperation and solidarity aimed at the conservation of biodiversity

(g) Enhancing the effectiveness, efficiency and transparency of resource provision and use;

**TARGET 20**

Strengthen **capacity-building** and development, access to and transfer of technology, and promote development of and access to innovation and technical and scientific **cooperation**, including through South- South, North-South and triangular cooperation, to meet the needs for effective implementation, particularly in developing countries, fostering joint technology development and joint scientific research programmes for the conservation and sustainable use of biodiversity and strengthening scientific research and monitoring capacities, commensurate with the ambition of the goals and targets of the framework.

**TARGET 21**

Ensure that the best available data, information and knowledge, are accessible to **decision makers**, **practitioners** and the public to guide effective and equitable governance, integrated and participatory management of biodiversity, and to strengthen communication, awareness-raising, education, monitoring, research and knowledge management and, also in this context, traditional knowledge, innovations, practices and technologies of indigenous peoples and local communities should only be accessed with their free, prior and informed consent20, in accordance with national legislation.

**TARGET 22**

Ensure the full, **equitable**, inclusive, effective and gender-responsive representation and participation in decision-making, and access to justice and information related to biodiversity by indigenous peoples and local communities, respecting their cultures and their rights over lands, territories, resources, and traditional knowledge, as well as by women and girls, children and youth, and persons with disabilities and ensure the full protection of environmental human rights defenders.

**TARGET 23**

Ensure gender **equality** in the **implementation** of the framework through a gender-responsive approach where all women and girls have equal opportunity and **capacity** to contribute to the three objectives of the Convention, including by recognizing their equal rights and access to land and natural resources and their

full, equitable, meaningful and informed participation and leadership at all levels of action, engagement, policy and decision-making related to biodiversity

**Table S1.2. Classification of keywords of 4 goals and 23 targets of the Kunming-Montreal global biodiversity framework (GBF) (CBD, 2020) and the monitoring elements from the zero draft of the GBF in terms of DPSIR classes (****driver, pressure, status, impact and response)** **(Burkhard and Müller, 2008), monitoring type (remotely observable characteristics, genetic resources, management policies) and habitat (aquatic, terrestrial, agnostic).**

| Classes | | Keywords | Included |
| --- | --- | --- | --- |
| **Driver** | | agriculture, harvest, quota, trade (of species), access | No |
| **Pressure** | | conflicts, (human induced) extinction, pollution, risk, invasive (species), acidification, densely populated | No |
| **State** | | Area’s (of high biodiversity importance/particular importance), connectivity, priority sites, extent, quality, integrity, connectivity, resilience, ecosystem function | Yes |
| **Impact** | | (nature’s) contributions to human, impacts, well being, ecosystem service, monetary, utilization, sustainable, consumption | No |
| **Response** | | Adaptation, administrative, control, conserved, eradication (of invasive species), implementation, enhanced, legal, maintained, management, measures, mitigation, practices, policy, recovery, restoration, strategies, sustainable, fair, equitable, capacity building, cooperation | No |
| Classes | | Keywords | Included |
| *Monitoring type* | |  |  |
| Remotely observable characteristics | | area, aquatic, degradation, extent, fauna, flora, fragmentation, marine, plants, quantity, quality, rate of change, reduction, species, terrestrial, wilderness | Yes |
| Genetic resources | | biotechnology, genetic, breed | No |
| Management policies | | accounts, awareness, capacity-building, consumption, cooperation, economy, education, financial resources, funding, knowledge, participation, policies, private, public, regulations, rights, strategy, subsidies, technology transfer, women, youth, planning levels | No |
| *Habitat* | |  |  |
| Aquatic | | aquaculture, aquatic, coastal, coral reefs, fish, fisheries, fresh water, mangroves, marine, sea grass, wetlands, | No |
| Terrestrial | | agriculture, farmland, forest, grassland, islands, land, terrestrial, wilderness, urban | Yes |
| Agnostic | | alien invasive species, climate change mitigation, natural ecosystems | Yes |

The identification of the relevant goals and targets of the final Kunming-Montreal global biodiversity framework was consistent with a previous analysis of the monitoring elements from the zero draft of the GBF. Our initial analysis focused on the monitoring elements from the zero draft as these descriptions better defined the focus for the different components of the targets and goals (see Table S1.2 in Supplementary Material S1). For this, we first filtered by the monitoring type, then by habitat type, and finally by DPSIR classes. Out of a total of 162, we identified 109 remotely observable monitoring elements (Figure S1.1), excluding those that were associated with ‘genetic resources’ (*n* = 6) and ‘management policies’ (*n* = 47). Of the 109 elements, 23 focused specifically on aquatic habitats and were subsequently excluded. A total of 86 elements focused either on terrestrial habitats (*n* = 45) or were agnostic (*n* = 41), and hence included. Out of these 86 elements, we identified 33 elements (using the keywords highlighted in Table S1.2. in Supplementary Materials S1) which can track the state of terrestrial biodiversity rather than pressures, drivers, responses or impacts (see Figure S1.1). By identifying the differences between the zero draft, 1^st^ draft and final text (given in Figure S2.1 and Table S2.1 of Supplementary Material S2), we found that the set of goals and targets included in our analysis was consistent, except for T12 (i.e., T11 in the updated zero draft version) which was only included when analysing the final text.

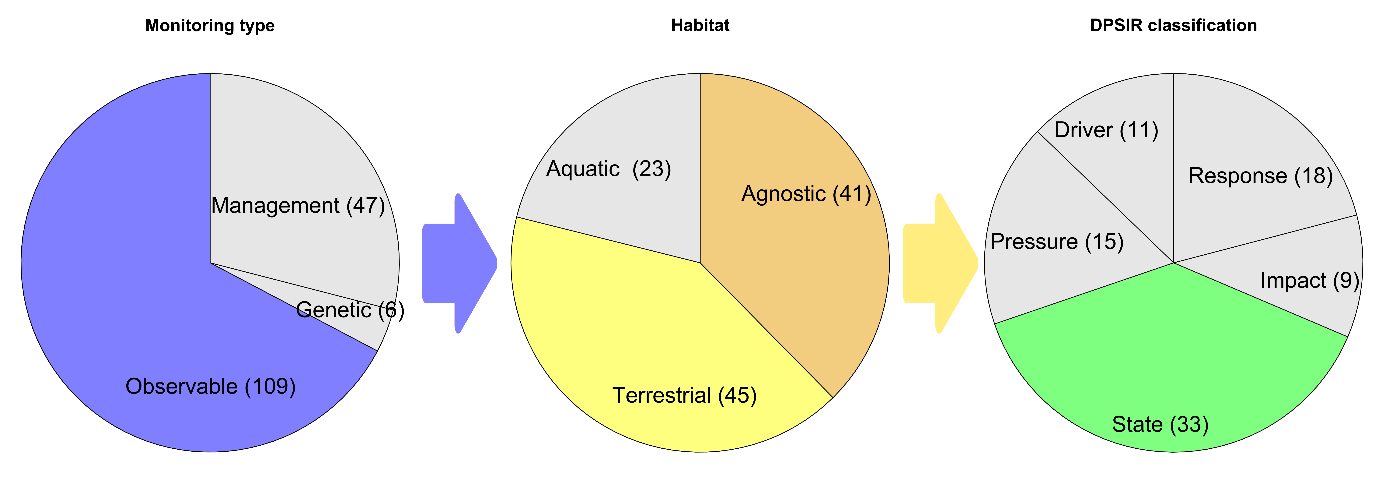

Figure S1.1. Number and proportion of monitoring elements at different stages of the filtering process (as described in Table S1.2. in Supplementary Materials S1). Included were only monitoring elements that are remotely observable with current instruments and technologies for terrestrial/agnostic habitats, and with a focus on the state of terrestrial biodiversity. DPSIR = driver, pressure, state, impact and response.

Table S1.3: Overview of all goals, targets, target components and monitoring elements for tracking the state of terrestrial biodiversity as described in the zero draft of the Kunming-Montreal global biodiversity framework of the Convention on Biological Diversity (CBD, 2020). Both monitoring elements and target components are no longer used in the final text of the Kunming-Montreal global biodiversity framework, and targets were adjusted accordingly (see modifications shown in Figure S2.1 of Supplementary Material S2). We use the descriptions as provided by the zero draft, including terms such as ‘other effective conservation measures’ (OECMs), ‘protected areas’ (PAs), and ‘species protection index’ (SPI). While target 11 was not identified on the basis of the description of its monitoring elements, it was added here to be consistent with the findings of the analysis of the final GBF text (see Table 3 of the main text)

| Goal | Target | Target component | Monitoring element |
| --- | --- | --- | --- |
| Goal A:  Reducing threats to biodiversity | T1: Ensure that [50%] of land and sea areas globally are under spatial planning addressing land/sea use change, retaining most of the existing intact and wilderness areas, and allow to restore [X%] of degraded freshwater, marine and terrestrial natural ecosystems and connectivity among them. | T1.1: Increase in area of terrestrial, freshwater and marine ecosystems under spatial planning | Trends in area under spatial land-use plans |
|  |  | T1.2: Prevention of reduction and fragmentation of natural habitats due to land/sea use change | Trends in extent and rate of change of forest ecosystems |
|  |  |  | Trends in extent and rate of change of dry and sub-humid lands |
|  |  |  | Trends in extent and rate of change of in grasslands |
|  |  |  | Trends in extent and rate of change of other terrestrial ecosystems |
|  |  |  | Trends in forest and agriculture lands as a proportion of total land area |
|  |  |  | Trends in farmland biodiversity and sustainability of agricultural land |
|  |  | T1.3: Priority retention of intact / wilderness areas | Trends in extent of natural intact/ wilderness |
|  |  | T1.4: Restoration of degraded ecosystems | Trends in the area of degraded forest ecosystems restored |
|  |  |  | Trends in the area of degraded dry and sub-humid lands restored |
|  |  |  | Trends in the area of degraded grassland ecosystems restored |
|  |  |  | Trends in the area of degraded other terrestrial ecosystems restored |
|  |  |  | Trend in the area of converted agricultural lands restored |
|  |  | T1.5: Maintenance and restoration of connectivity of natural ecosystems | Trends in habitat connectivity |
|  | T2: Protect and conserve through well connected and effective system of protected areas and other effective area-based conservation measures at least 30 per cent of the planet with the focus on areas particularly important for biodiversity. | T2.1: Area of terrestrial, freshwater and marine ecosystem under protection and conservation | Trends in extent of protected areas |
|  |  |  | Trends in extent of areas under other area-based conservation measures |
|  |  | T2.2: Areas of particular importance for biodiversity are protected and conserved as priority | Trends in proportion of areas of particular importance for biodiversity protected and conserved |
|  |  | T2.3: Representative system of protected areas and other effective area-based conservation measures | Trends in ecological representativeness of areas conserved |
|  |  | T2.5: Effective and equitable management of the system of protected areas and other effective area-based conservation measures | Trends in connectivity of protected areas and other effective area-based conservation measures |
|  |  | T2.6: Increased protection and conservation effectiveness | Trends in conservation effectiveness of protected areas and other area-based conservation measures |
|  |  | T2.7: Integration into landscape and seascape context | Trends in policy and governance practices outside of protected areas and OECMs compatible with their trends in management objectives |
|  | T5: Manage, and where possible control, pathways for the introduction of invasive alien species, achieving [50%] reduction in the rate of new introductions, and control or eradicate invasive alien species to eliminate or reduce their impacts, including in at least [50%] of priority sites. | T5.2: Effective detection, identification, prioritisation and monitoring of invasive alien species | Trends in efficiency of detection of invasive alien species |
|  |  |  | Trends in identification of invasive alien species |
|  |  |  | Trends in monitoring of invasive alien species |
|  | T7: Increase contributions to climate change mitigation adaption and disaster risk reduction from nature-based solutions and ecosystems based approaches, ensuring resilience and minimizing any negative impacts on biodiversity | T7.1: Increased biodiversity contribution to climate change mitigation, adaptation and disaster risk reduction | Trends in carbon stocks in different ecosystems |
|  |  |  | Trends in contribution to climate change adaptation |
|  |  |  | Trends in contribution to disaster risk reduction |
|  |  |  | Trends in invertebrate stocks |
| Goal B:  Meeting people’s needs | T9: Support the productivity, sustainability and resilience of biodiversity in agricultural and other managed ecosystems through conservation and sustainable use of such ecosystems, reducing productivity gaps by at least [50%]. | T9.1: Sustainable management of agricultural biodiversity, including soil biodiversity, cultivated plants and farmed and domesticated animals and of wild relatives | Trends in soil quality |
|  |  |  | Trends in pollinators |
|  |  |  | Trends in genetic diversity of cultivated plants and of wild relatives |
|  |  |  | Trends in genetic diversity of domesticated animals and of wild relatives |
|  | T10: Ensure that, nature based solutions and ecosystem approach contribute to regulation of air quality, hazards and extreme events and quality and quantity of water for at least [XXX million] people. | T10.1: Regulation of air quality | Trends in extent and quality of natural ecosystems |
|  |  | T10.2: Regulation of freshwater quantity, quality, location and timing | Trends in extent and quality of natural ecosystems |
|  |  | T10.3: Regulation of hazards and extreme events | Trends in extent and quality of natural ecosystems |
|  | T11. By 2030, increase benefits from biodiversity and green/blue spaces for human health and well-being, including the proportion of people with access to such spaces by at least [100%], especially for urban dwellers | T11.1 Access to green/blue spaces | Trends in access to green/blue spaces |
|  |  | T11.2 Contributions of biodiversity to human health and well-being | Trends in species that provide essential services  Trends in contributions to human health and well-being from forest ecosystems  Trends in contributions to human health and well-being from dry and sub-humid lands  Trends in contributions to human health and well-being from grasslands  Trends in contributions to human health and well-being from other terrestrial ecosystems  Trends in contributions to human health and well-being from mangroves  Trends in contributions to human health and well-being from coral reefs  Trends in contributions to human health and well-being from seagrass ecosystems  Trends in contributions to human health and well-being from coral reefs  Trends in contributions to human health and well-being from wetlands |
