## Supplementary Materials S2 for "Advancing terrestrial biodiversity monitoring with satellite remote sensing in the context of the Kunming-Montreal global biodiversity framework"

### Supplementary Material S2

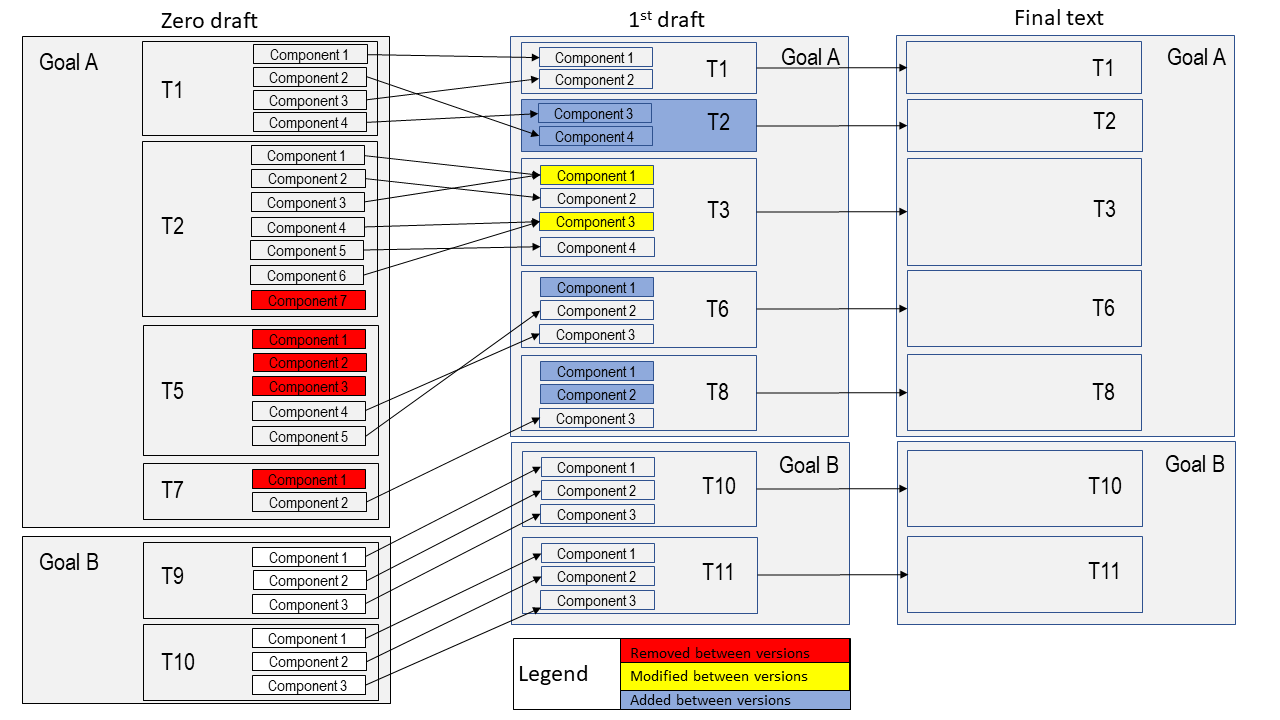

Figure S2.1: Structural changes between the zero draft (CBD, 2020), the 1^st^ draft (CBD, 2021) and the final text (CBD, 2022a, 2022b) of the Kunming-Montreal global biodiversity framework. Shown are only those targets and target components that are associated with monitoring elements which focus on the state of terrestrial biodiversity and which are observable from space (our focus, see methods in main text). The colouring of the cells indicates whether a component has been removed (red), significantly modified through merging of multiple components into one (yellow) or added as a new component (blue). See Table S2.1 for details of the changes.

Table S2.1: Details of the textual changes in the global biodiversity framework from the zero draft (left) to the 1^st^ draft (middle) and the final version (right). The highlighted text (i.e. colouring of the cells) indicates whether a component has been removed (red), significantly modified through merging of multiple components into one (yellow) or added as new component (blue). Small textual changes are highlighted as moderate changes (brown font underlined) or added text (blue font). We use the descriptions as provided by the two drafts, including terms such as ‘GtCO2e’ referring to ‘Gigatons of emitted CO2’.

| Goal | Target | Component |  | Component | Target |  | Target |
| --- | --- | --- | --- | --- | --- | --- | --- |
| Goal A: The integrity, connectivity and resilience of all ecosystems are maintained, enhanced, or restored, substantially increasing the area of natural ecosystems by 2050; Human induced extinction of known threatened species is halted, and, by 2050, extinction rate and risk of all species are reduced tenfold and the abundance of native wild species is increased to healthy and resilient levels; The genetic diversity within populations of wild and domesticated species, is maintained, safeguarding their adaptive potential. | T1: Ensure that [50%] of land and sea areas globally are under spatial planning addressing land/sea use change, retaining most of the existing intact and wilderness areas, and   allow to restore [X%] of degraded freshwater, marine and terrestrial natural ecosystems and connectivity among them. | T1.1: Increase in area of terrestrial, freshwater and marine ecosystems under spatial planning | > | T1.1: Area under integrated biodiversity-inclusive spatial planning | T1. Ensure that all land and sea areas globally are under integrated biodiversity-inclusive spatial planning addressing land- and sea-use change, retaining existing intact and wilderness areas. | > | T1. Ensure that all areas are under participatory integrated biodiversity inclusive spatial planning and/or effective management processes addressing land and sea use change, to bring the loss of areas of high biodiversity importance, including ecosystems of high ecological integrity, close to zero by 2030, while respecting the rights of indigenous peoples and local communities |
|  |  | T1.3: Priority retention of intact / wilderness areas | > | T1.2: Retention of existing intact and wilderness areas |  | > |  |
|  |  | T1.4: Restoration of degraded ecosystems | > | T2.1: Area of freshwater, marine and terrestrial ecosystems restored | T2: Ensure that at least 20 percent of degraded freshwater, marine and terrestrial ecosystems are under restoration, ensuring connectivity among them and focusing on priority ecosystems. | > | T2. Ensure that by 2030 at least 30 per cent of areas of degraded terrestrial, inland water, and coastal and marine ecosystems are under effective restoration, in order to enhance biodiversity and ecosystem functions and services, ecological integrity and connectivity. |
|  |  | T1.2: Prevention of reduction and fragmentation of natural habitats due to land / sea use change | > | T2.2: Connectivity |  | > |  |
|  | T2: Protect and conserve through well connected and effective system of protected areas and other effective area-based conservation measures at least 30 per cent of the planet with the focus on areas particularly important for biodiversity. | T2.1: Area of terrestrial, freshwater and marine ecosystem under protection and conservation | > | T3.1: Area protected and conserved | T3: Ensure that at least 30 per cent globally of land areas and of sea areas, especially areas of particular importance for biodiversity and its contributions to people, are conserved through effectively and equitably managed, ecologically representative and well-connected systems of protected areas and other effective area-based conservation measures and integrated into the wider landscapes and seascapes. | > | T3. Ensure and enable that by 2030 at least 30 per cent of terrestrial, inland water, and of coastal and marine areas, especially areas of particular importance for biodiversity and ecosystem functions and services, are effectively conserved and  managed through ecologically representative, well-connected and equitably governed systems of protected areas and other effective area-based conservation measures, recognizing indigenous and traditional territories, where applicable,  and integrated into wider landscapes, seascapes and the ocean, while ensuring that any sustainable use, where appropriate in such areas, is fully consistent with conservation outcomes, recognizing and respecting the rights of indigenous peoples and local communities including over their traditional territories, |
|  |  | T2.3: Representative system of protected areas and other effective area-based conservation measures | > |  |  | > |  |
|  |  | T2.2: Areas of particular importance for biodiversity are protected and conserved as priority | > | T3.2: Areas of particular importance for biodiversity protected and conserved |  | > |  |
|  |  | T2.4: Effective and equitable management of the system of protected areas and other effective area-based conservation measures | > | T3.3: Effective management and equitable governance of the system of protected areas and other effective area-based conservation measures |  | > |  |
|  |  | T2.6: Increased protection and conservation effectiveness | > |  |  | > |  |
|  |  | T2.5: Connectivity within the system of protected areas and other effective area-based conservation measures |  | T3.4: Connectivity within the system of protected areas and other effective area-based conservation measures |  |  |  |
|  |  | T2.7: Integration into landscape and seascape context |  | - |  |  |  |
|  | T5: Manage, and where possible control, pathways for the introduction of invasive alien species, achieving [50%] reduction in the rate of new introductions, and control or eradicate invasive alien species to eliminate or reduce their impacts, including in at least [50%] of priority sites. | T5.1: Management of pathways for introduction of invasive alien species | > |  | T6: Manage pathways for the introduction of invasive alien species, preventing, or reducing their rate of introduction and establishment by at least 50 per cent, and control or eradicate invasive alien species to eliminate or reduce their impacts, focusing on priority species and priority sites. | > | T6. Eliminate, minimize, reduce and or mitigate the impacts of invasive alien species on biodiversity and ecosystem services by identifying and managing pathways of the introduction of alien species, preventing the introduction and establishment of priority invasive alien species, reducing the rates of introduction and establishment of other known or potential invasive alien species by at least 50 per cent, by 2030, eradicating or controlling invasive alien species especially in priority sites, such as islands . |
|  |  | T5.2: Effective detection, identification, prioritisation and monitoring of invasive alien species |  |  |  |  |  |
|  |  | T5.3: Establishment of measures for eradication, control and management of invasive alien species |  |  |  |  |  |
|  |  | T5.4: Eliminated or reduced impacts of IAS | > | T6.3: Reducing the impact on priority species and priority sites |  | > |  |
|  |  | T5.5: Eradication, control or management of IAS in priority sites | > | T6.2: Control or eradicate invasive alien species |  | > |  |
|  |  |  |  | T6.1: Rate of introduction and establishment |  |  |  |
|  | T7: Increase contributions to climate change mitigation adaption and disaster risk reduction from nature-based solutions and ecosystems based approaches, ensuring resilience and minimizing any negative impacts on biodiversity |  |  | T8.1: Minimize impact of climate change | T8: Minimize the impact of climate change on biodiversity, contribute to mitigation and adaptation through ecosystem-based approaches, contributing at least 10 GtCO2e per year to global mitigation efforts, and ensure that all mitigation and adaptation efforts avoid negative impacts on biodiversity |  | T8. Minimize the impact of climate change and ocean acidification on biodiversity and increase its resilience through mitigation, adaptation, and disaster risk reduction actions, including through nature-based solution and/or ecosystem based approaches, while minimizing negative and fostering positive impacts of climate action on biodiversity. |
|  |  |  |  | T8.2: Contribute at least 10 GtCO2 to mitigation and adaptation through ecosystem-based approaches |  |  |  |
|  |  | T7.2: Minimised negative impacts on biodiversity from any mitigation, adaptation and disaster risk reduction measures |  | T8.3: Ensure that all mitigation and adaptation efforts avoid negative impacts on biodiversity |  |  |  |
|  |  | T7.1: Increased biodiversity contribution to climate change mitigation, adaptation and disaster risk reduction |  | - |  |  |  |
| Goal B: Biodiversity is sustainably used and managed and nature’s contributions to people, including ecosystem functions and services, are valued, maintained and enhanced, with those currently in decline being restored, supporting the achievement of sustainable development for the benefit of present and future generations by 2050. | T9: Support the productivity, sustainability and resilience of biodiversity in agricultural and other managed ecosystems through conservation and sustainable use of such ecosystems, reducing productivity gaps by at least [50%]. | T9.1: Sustainable management of agricultural biodiversity, including soil biodiversity, cultivated plants and farmed and domesticated animals and of wild relatives | > | T10.1: Agriculture | T10: Ensure all areas under agriculture, aquaculture and forestry are managed sustainably, in particular through the conservation and sustainable use of biodiversity, increasing the productivity and resilience of these production systems. | > | T10. Ensure that areas under agriculture, aquaculture, fisheries and forestry are managed sustainably, in particular through the sustainable use of biodiversity, including through a substantial increase of the application of biodiversity friendly practices, such as sustainable intensification, agroecological and other innovative approaches contributing to the resilience and long-term efficiency and productivity of these production systems and to food security, conserving and restoring biodiversity and maintaining nature’s contributions to people, including ecosystem functions and services |
|  |  | T9.2: Sustainable management of aquaculture | > | T10.2: Aquaculture |  | > |  |
|  |  | T9.3: Sustainable management of all types of forests | > | T10.3: Forestry |  | > |  |
|  | T10: Ensure1 that, nature based solutions and ecosystem approach contribute to regulation of air quality, hazards and extreme events and quality and quantity of water for at least [XXX million] people. | T10.1 Regulation of air quality | > | T11.1: Air quality | T11: Maintain and enhance nature’s contributions to regulation of air quality, quality and quantity of water, and protection from hazards and extreme events for all people. | > | T11. Restore, maintain and enhance nature’s contributions to people, including ecosystem functions and services, such as regulation of air, water, and climate, soil health, pollination and reduction of disease risk, as well as protection from natural hazards and disasters, through nature-based solutions and ecosystem based approaches for the benefit of all people and nature. |
|  |  | T10.3 Regulation of freshwater quantity, quality, location and timing | > | T11.2: Quality and quantity of water |  | > |  |
|  |  | T10.2: Regulation of hazards and extreme events | > | T11.3: Protection from hazards and extreme events |  | > |  |
|  | T11: By 2030, increase benefits from biodiversity and green/blue spaces for human health and well-being, including the proportion of people with access to such spaces by at least [100%], especially for urban dwellers. | - |  | T12.1: Increase the area of green and blue spaces | T12: Increase the area of, access to, and benefits from green and blue spaces, for human health and well-being in urban areas and other densely populated areas |  | T12: Significantly increase the area and quality and connectivity of, access to, and benefits from green and blue spaces in urban and densely populated areas sustainably, by mainstreaming the conservation and sustainable use of biodiversity, and ensure biodiversity-inclusive urban planning, enhancing native biodiversity, ecological connectivity and integrity, and improving human health and well-being and connection to nature and contributing to inclusive and sustainable urbanization and the provision of ecosystem functions and  services. |
|  |  | T11.1  Access to green/blue spaces |  | T12.2: Increase the access to, and benefits from, green and blue spaces |  |  |  |
|  |  | T11.2  Contributions of biodiversity to human health, and well-being |  | T12.3 Increase the access to and benefits from green and blue spaces |  |  |  |
