## Supplementary Materials S3 for "Advancing terrestrial biodiversity monitoring with satellite remote sensing in the context of the Kunming-Montreal global biodiversity framework"

### Supplementary Material S3

Table S3.1: Biodiversity indicators associated with the goals and targets of the Kunming-Montreal global biodiversity framework (CBD, 2022). The list represents all headline, component and complementary indicators for tracking the state of terrestrial biodiversity (represented in the ID column with h, c, and aux, respectively). The colouring of the cells indicates whether a biodiversity indicator is eligible for tracking the state of terrestrial biodiversity using spatial information products (blue = TRUE, orange = FALSE), and whether an eligible indicator is considered mature (blue = TRUE, orange = FALSE, brown = in development). The maturity of non-eligible indicators was not explored and therefore left blank.

| Indicators | ID | Eligible | Mature |
| --- | --- | --- | --- |
| Above-ground biomass stock in forest (tonnes/ha) | T08.aux.1 | 1 | 1 |
| Agrobiodiversity Index | T10.aux.1 | 1 | 1 |
| Air emission accounts | T11.aux.1 | 0 |  |
| Air pollution emissions account | B.aux.8 | 0 |  |
| Air quality index | B.aux.7 | 0 |  |
| Annual mean levels of fine particulate matter (e.g., PM2.5 and PM10) in cities | T11.c.3 | 0 |  |
| Annual Tropical Primary Tree Cover Loss | T02.aux.4 | 1 | 1 |
| Area of forest under sustainable management: total forest management certification by Forest Stewardship Council and Programme for the Endorsement of Forest Certification | T10.c.1 | 1 | 1 |
| Area under restoration | T02.h.1 | 0 |  |
| Average income of small-scale food producers, by sex and indigenous status | T10.c.2 | 0 |  |
| Average share of the built-up area of cities that is green/blue space for public use for all | T12.h.1 | 1 | 1 |
| Bioclimatic Ecosystem Resilience Index (BERI) | A.aux.27 | 1 | 1 |
| Bioclimatic Ecosystem Resilience Index (BERI) | T02.aux.9 | 1 | 1 |
| Bioclimatic Ecosystem Resilience Index (BERI) | T08.c.4 | 1 | 1 |
| Biodiversity Habitat Index | A.c.4 | 1 | 1 |
| Biodiversity Habitat Index | A.aux.25 | 1 | 1 |
| Biodiversity Habitat Index | T02.aux.12 | 1 | 1 |
| Biodiversity Intactness Index | A.aux.30 | 1 | 1 |
| Biofuel production | B.aux.20 | 0 |  |
| Biomass of selected natural ecosystems | A.aux.24 | 1 | - |
| Carbon stocks and annual net GHG emissions, by landuse category, split by natural and non-natural land cover | T08.aux.6 | 0 |  |
| Change in the extent of inland water ecosystems over time | A.aux.21 | 0 |  |
| Change in the extent of water-related ecosystems over time | A.c.9 | 0 |  |
| Change in the quality of coastal water ecosystems over time | B.aux.16 | 0 |  |
| Change in the quality of inland water ecosystems over time | B.aux.15 | 0 |  |
| Changes in plankton biomass and abundance | A.aux.41 | 0 |  |
| Changes in soil organic carbon stocks | T10.aux.2 | 1 | 1 |
| Climatic impact index | B.aux.10 | 1 | 1 |
| CMS Connectivity Indicator | A.aux.50 | 1 | - |
| CMS Connectivity Indicator | T03.c.6 | 1 | - |
| Comprehensiveness of conservation of socioeconomically as well as culturally valuable species | A.aux.46 | 0 |  |
| Conservation (proposed binary indicator to be further developed) | T01.aux.10 | 0 | 0 |
| Continuous Global Mangrove Forest Cover | A.aux.8 | 0 | 0 |
| Cover of key benthic groups | A.aux.18 | 0 |  |
| Coverage of protected areas and OECMs (3.1) | T03.h.1 | 0 |  |
| Coverage of Protected areas and OECMS and traditional territories (by governance type) | T03.aux.6 | 0 |  |
| Cultural vitality index | B.aux.31 | 0 |  |
| Dendritic Connectivity Index | A.aux.35 | 0 |  |
| Ecological Footprint | B.aux.2 | 0 |  |
| Ecosystem Intactness Index | A.c.1 | 1 | 1 |
| Ecosystem Intactness Index | A.aux.29 | 1 | 1 |
| Ecosystem Integrity Index | A.c.2 | 1 | 1 |
| Ecosystem Integrity Index | T01.aux.11 | 1 | 1 |
| EDGE | A.c.7 | 0 |  |
| Eflow index | B.aux.14 | 0 |  |
| Expected Loss of Phylogenetic diversity | A.aux.53 | 0 |  |
| Expected loss of Phylogenetic Diversity | B.aux.4 | 0 |  |
| Extent of indigenous peoples and local communities’ lands that have some form of recognition | T03.aux.9 | 0 |  |
| Extent of natural ecosystems | A.h.2 | 1 | - |
| Extent of natural ecosystems | T01.h.2 | 1 | - |
| Extent of natural ecosystems by type | T01.aux.8 | 1 | - |
| Extent of natural ecosystems by type | T02.c.1 | 1 | - |
| Extent of physical damage indicator to predominant seafloor habitats physical damage | A.aux.32 | 0 |  |
| Extent to which protected areas and other effective area based conservation measures (OECMs) cover Key Biodiversity Areas that are important for migratory species | T03.aux.5 | 0 |  |
| Fish abundance and biomass | A.aux.42 | 0 |  |
| Fleshy algae cover | A.aux.19 | 0 |  |
| Forest area as a proportion of total land area | A.aux.1 | 1 | 1 |
| Forest distribution | A.aux.2 | 1 | - |
| Forest Fragmentation Index | A.aux.22 | 1 | - |
| Forest Landscape Integrity Index | A.aux.23 | 1 | 1 |
| Forest Landscape Integrity Index | T02.aux.5 | 1 | 1 |
| Forestry Production & Trade (Wood Fuel) | B.aux.24 | 0 |  |
| Free flowing rivers | A.aux.55 | 0 |  |
| Free flowing rivers | T02.aux.7 | 0 |  |
| Genetic scorecard for wild species | A.aux.43 | 0 |  |
| Global coral reef extent | A.aux.13 | 0 |  |
| Global Ecosystem Restoration Index | T02.aux.6 | 1 | 1 |
| Global saltmarsh extent | A.aux.15 | 0 |  |
| Global Seagrass Extent (Seagrass Cover and composition) | A.aux.14 | 0 |  |
| Global Vegetation Health Products | A.aux.26 | 1 | 1 |
| Grassland and savannah extent | A.aux.4 | 1 | - |
| Green status index (pollinators) | B.aux.6 | 1 | - |
| Habitat distributional range | T02.aux.1 | 0 |  |
| Habitat patches located within marine protected areas or integrated coastal zone management (ICZM) | T01.aux.3 | 0 |  |
| Hard Coral cover and composition | A.aux.12 | 0 |  |
| Increase in secondary natural forest cover | T02.aux.3 | 1 | 0.5 |
| Index of coastal eutrophication | T08.aux.5 | 0 |  |
| Index of development of the standard - setting framework for the protection and promotion of culture, cultural rights and cultural diversity | B.aux.30 | 0 |  |
| Index of Linguistic Diversity - Trends of Bilinguistic diversity and numbers of speakers of indigenous languages | B.aux.29 | 0 |  |
| Index of Species Rarity Sites, High Biodiversity Areas, Large Mammal Landscapes, Intact Wilderness and Climate Stabilization Areas | T02.aux.2 | 1 | 1 |
| Intact Wilderness | A.aux.52 | 1 | - |
| Intact Wilderness | B.aux.19 | 1 | - |
| Kelp canopy extent | A.aux.16 | 0 |  |
| Level of erosion | B.aux.17 | 0 |  |
| Level of water stress | T11.c.5 | 0 |  |
| Level of water stress: freshwater withdrawal as a proportion of available freshwater resources | B.aux.12 | 0 |  |
| Levels of poverty in biodiversity depended communities | B.aux.1 | 0 |  |
| Live coral cover | A.aux.11 | 0 |  |
| Living Planet Index | A.c.8 | 1 | 1 |
| Living Planet Index | B.c.2 | 1 | 1 |
| Living Planet Index | T02.aux.15 | 1 | 1 |
| Macroalgal Canopy Cover and Composition | A.aux.17 | 0 |  |
| Maintenance and restoration of connectivity of natural ecosystems | T02.c.2 | 0 |  |
| Marine species richness | A.aux.45 | 0 |  |
| Maximum fish catch potential | B.aux.21 | 0 |  |
| Mean Species Abundance (MSA) | A.aux.39 | 0 |  |
| Mortality rate attributed to household and ambient air pollution (SDG indicator 3.9.1) | T11.aux.4 | 0 |  |
| Mortality rate attributed to unsafe water, unsafe sanitation and lack of hygiene (exposure to unsafe Water, Sanitation and Hygiene for All (WASH) services | T11.c.2 | 0 |  |
| Mountain Green Cover Index | A.aux.5 | 1 | 1 |
| National greenhouse inventories from land use and land use change | T08.aux.2 | 0 |  |
| National greenhouse inventories from land use and land use change | T08.c.3 | 0 |  |
| Number of certified forest areas under sustainable management with verified impacts on biodiversity conservation | B.aux.3 | 1 | 1 |
| Number of countries adopting relevant national legislation and adequately resourcing the prevention or control of invasive alien species (proposed binary indicator to be further developed) | T06.aux.4 | 0 |  |
| Number of countries implementing national legislation, policies or other measures regarding FPIC related to conservation | T03.aux.11 | 0 |  |
| Number of countries implementing national legislation, policies or other measures regarding FPIC related to conservation” would work here for IPs (not necessarily LCs), if ‘spatial planning’ was substituted for | T01.aux.9 | 0 |  |
| Number of countries that adopt and implement national disaster risk reduction strategies in line with the Sendai Framework for Disaster Risk Reduction 2015–2030 which include biodiversity | T08.c.2 | 0 |  |
| Number of countries using natural capital accounts in planning processes | T01.aux.1 | 0 |  |
| Number of countries using ocean accounts in planning processes | T01.aux.5 | 0 |  |
| Number of countries with nationally determined contributions, long-term strategies, national adaptation plans and adaptation communications that reflect biodiversity (proposed binary indicator to be further developed) | T08.aux.7 | 0 |  |
| Number of deaths, missing persons and directly affected persons attributed to disasters per 100,000 population | B.aux.18 | 0 |  |
| Number of deaths, missing persons and directly affected persons attributed to disasters per 100,000 population | T11.c.1 | 0 |  |
| Number of formal and nonformal education programmes transmitting spiritual and cultural values in the UNESCO World Network of Biosphere Reserves | B.aux.27 | 0 |  |
| Number of hectares of UNESCO designated sites (natural and mixed World Heritage sites and Biosphere Reserves) | T03.aux.1 | 0 |  |
| Number of invasive alien species in national lists as per the Global Register of Introduced and Invasive Species | T06.aux.1 | 1 | 1 |
| Number of invasive alien species introduction events | T06.c.3 | 1 | 1 |
| Number of least developed countries and small island developing States with nationally determined contributions, long-term strategies, national adaptation plans, strategies as reported in adaptation communications and national communications | T08.aux.4 | 0 |  |
| Number of mixed sites (having both natural and cultural Outstanding Universal Values), cultural landscapes (recognized as combined works of nature and people) and natural sites with cultural values including those supporting local and indigenous knowledge and practices inscribed on the UNESCO World Heritage List and UNESCO World Network of Biosphere Reserves | B.aux.28 | 0 |  |
| Number of plant and animal genetic resources for food and agriculture secured in either medium- or long-term conservation facilities | A.aux.47 | 0 |  |
| Number of threatened species by species group | A.aux.37 | 0 |  |
| Ocean acidification | B.aux.11 | 0 |  |
| Ocean Health Index | A.aux.31 | 0 |  |
| Other spatial management plans (not captured as ICZM or marine spatial planning in 14.2.1) | T01.aux.4 | 0 |  |
| Parc connectedness | A.c.6 | 1 | 1 |
| Peatland extent and condition | A.aux.6 | 1 | - |
| Percent of land and seas covered by biodiversity-inclusive spatial plans * (1.1) | T01.h.3 | 1 | - |
| Percent of total land area that is under cultivation | T01.aux.7 | 1 | 1 |
| Percentage of biosphere reserves that have a positive conservation outcome and effective management | T03.aux.8 | 0 |  |
| Percentage of cropped landscapes with at least 10 % natural land | T02.aux.8 | 1 | - |
| Percentage of spatial plans utilizing information on key biodiversity areas | T01.aux.2 | 0 |  |
| Percentage of threatened species that are improving in status according to the Red List | A.aux.36 | 1 | - |
| Permafrost thickness, depth and extent | A.aux.7 | 0 |  |
| Population involved in hunting and gathering | B.aux.22 | 0 |  |
| Prevalence of moderate or severe food insecurity in the population, based on the Food Insecurity Experience Scale | B.aux.23 | 0 |  |
| Priority retention of intact / wilderness areas | T02.aux.10 | 0 |  |
| Priority retention of intact/ wilderness areas | T01.c.1 | 0 |  |
| Processes and tools to monitor the implementation of a right to a healthy environment (e.g. Included in NBSAPs and reported in national reports | B.aux.33 | 0 |  |
| Progress towards sustainable forest management (10.2) | T10.h.2 | 1 | 0.5 |
| Proportion of agricultural area under productive and sustainable agriculture (10.1) | T10.h.1 | 1 | - |
| Proportion of bodies of water with good ambient water quality | B.aux.13 | 0 |  |
| Proportion of bodies of water with good ambient water quality | T11.c.4 | 0 |  |
| Proportion of land that is degraded over total land area | T10.aux.6 | 1 | 1 |
| Proportion of local administrative units with established and operational policies and procedures for participation of local communities in water and sanitation management | T11.aux.2 | 0 |  |
| Proportion of local breeds classified as being at risk of extinction | T10.aux.5 | 0 |  |
| Proportion of local breeds classified as being at risk, extinction | A.aux.48 | 0 |  |
| Proportion of local governments that adopt and implement local disaster risk reduction strategies in line with national disaster risk reduction strategies | T08.aux.3 | 0 |  |
| Proportion of population using safely managed drinking water services | T11.aux.3 | 0 |  |
| Proportion of populations maintained within species | A.aux.54 | 0 |  |
| Proportion of terrestrial, freshwater and marine ecological regions which are conserved by protected areas or other effective area-based conservation measures | T03.aux.13 | 1 | 1 |
| Proportion of transboundary basin area with an operational arrangement for water cooperation | T01.aux.6 | 0 |  |
| Protected area and OECM management effectiveness (MEPCA) indicator | T03.aux.2 | 0 |  |
| Protected Area Connectedness Index (PARC-Connectedness) | T03.c.4 | 1 | 1 |
| Protected area coverage of key biodiversity areas | T03.c.1 | 0 |  |
| Protected Area Isolation Index (PAI) | T03.aux.3 | 1 | 1 |
| Protected Area Management Effectiveness (PAME) | T03.c.2 | 0 |  |
| Protected Areas Network metric (ProNet) | T03.aux.4 | 0 |  |
| Protected Connected (Protconn) index | A.c.5 | 1 | 1 |
| Protected Connected (Protconn) index | T03.c.3 | 1 | 1 |
| Ramsar Management Effectiveness Tracking Tool (RMETT) | T03.aux.7 | 0 |  |
| Rate of invasive alien species establishment (6.1) | T06.h.1 | 1 | - |
| Rate of invasive alien species spread | T06.c.2 | 1 | 1 |
| Rate of invasive species impact and rate of impact | T06.c.1 | 1 | - |
| Recreation and cultural ecosystem services provided | T12.c.1 | 0 |  |
| Red List Index | A.h.3 | 1 | 1 |
| Red List Index | A.aux.49 | 1 | 1 |
| Red List Index | B.c.1 | 1 | 1 |
| Red List Index | B.aux.5 | 1 | 1 |
| Red List Index | B.aux.34 | 1 | 1 |
| Red List Index | T02.aux.13 | 1 | 1 |
| Red List Index | T06.aux.3 | 1 | 1 |
| Red List Index | T10.aux.3 | 1 | 1 |
| Red List Index | T10.aux.4 | 1 | 1 |
| Red List of Ecosystems | A.h.1 | 1 | 1 |
| Red List of Ecosystems | T01.h.1 | 1 | 1 |
| Red List of Ecosystems | T02.aux.14 | 1 | 1 |
| Red List of Ecosystems | T03.c.5 | 1 | 1 |
| Red List of Ecosystems | T03.aux.12 | 1 | 1 |
| Relative Magnitude of Fragmentation (RMF) | A.aux.28 | 0 |  |
| River Fragmentation Index | A.aux.34 | 0 |  |
| Services provided by ecosystems* (B.1) | B.h.1 | 0 |  |
| Services provided by ecosystems* (B.1) | T11.h.1 | 0 |  |
| Species habitat Index | A.c.3 | 1 | 1 |
| Species habitat Index | T02.aux.16 | 1 | 1 |
| Species Protection Index | A.aux.40 | 1 | 1 |
| Species Protection Index | T03.c.8 | 1 | 1 |
| Species Protection Index | T03.aux.10 | 1 | 1 |
| Species richness/Changes in local terrestrial diversity (PREDICTS) | A.aux.44 | 1 | 1 |
| Species Status Index | A.aux.51 | 1 | - |
| Status of Key Biodiversity Areas | T02.aux.11 | 1 | - |
| The number of protected areas that have completed a site-level assessment of governance and equity (SAGE) | T03.c.7 | 0 | 0 |
| The proportion of populations within species with an effective population size > 500 (A.5) | A.h.5 | 0 | 0 |
| Total climate regulation services provided by ecosystems by ecosystem type (System of Environmental Economic Accounts) | T08.c.1 | 1 | - |
| Tree cover loss | A.aux.3 | 1 | 1 |
| Trends in abundance, temporal occurrence, and spatial distribution of non-indigenous species, particularly invasive, non-indigenous species, notably in risk areas (in relation to the main vectors and pathways of spreading of such species) | T06.aux.2 | 0 |  |
| Trends in mangrove extent | A.aux.10 | 0 |  |
| Trends in mangrove forest fragmentation | A.aux.9 | 0 |  |
| Trends in the legal trade of medicinal plants | B.aux.25 | 0 |  |
| UNESCO Culture 2030 (multiple indicators) | B.aux.32 | 0 |  |
| Visitor management assessment | B.aux.26 | 0 |  |
| Wetland Extent Trends Index | A.aux.20 | 0 |  |
| Wetland Extent Trends Index | A.aux.33 | 0 |  |
| Wild bird index | A.aux.38 | 1 | 1 |
| Zoonotic disease in wildlife | B.aux.9 | 0 |  |
| Total |  | 86 |  |
| Total Unique |  | 57 | 35 mature |

Table S3.2: Biodiversity indicators associated with the Kunming-Montreal global biodiversity framework (CBD, 2022) which were included in the workflow analysis, with sources of the workflow description.

| Nr | Biodiversity indicators | Source | Count |
| --- | --- | --- | --- |
| 1 | Above-ground biomass stock in forest (tonnes/ha) | (FAO, 2020) | 1 |
| 2 | Agrobiodiversity Index | (Jones et al., 2021) | 1 |
| 3 | Annual Tropical Primary Tree Cover Loss | (Global Forest Watch, 2018) | 1 |
| 4 | Area of forest under sustainable management: total forest management certification by Forest Stewardship Council and Programme for the Endorsement of Forest Certification | (Biodiversity Indicators Partnership, 2013) | 1 |
| 5 | Average share of the built-up area of cities that is green/blue space for public use for all | (UN-Habitat, 2018) | 1 |
| 6 | Bioclimatic Ecosystem Resilience Index (BERI) | (Ferrier et al., 2020, 2007) | 3 |
| 7 | Biodiversity Habitat Index | (Hoskins et al., 2016) | 3 |
| 8 | Biodiversity Intactness Index | (Newbold et al., 2016) | 1 |
| 9 | Changes in soil organic carbon stocks | (Lorenz et al., 2019) | 1 |
| 10 | Climatic impact index | (Stephens et al., 2016) | 1 |
| 11 | Ecosystem Intactness Index | (Beyer et al., 2020) | 2 |
| 12 | Ecosystem Integrity Index | (Hill et al., 2022) | 2 |
| 13 | Forest area as a proportion of total land area | (FAO, 2020) | 1 |
| 14 | Forest Landscape Integrity Index | (Grantham et al., 2020) | 2 |
| 15 | Global Ecosystem Restoration Index | (Fernandez et al., 2015) | 1 |
| 16 | Global Vegetation Health Products | (Kogan, 1997) | 1 |
| 17 | Index of Species Rarity Sites, High Biodiversity Areas, Large Mammal Landscapes, Intact Wilderness and Climate Stabilization Areas | (Dinerstein et al., 2020) | 1 |
| 18 | Living Planet Index | (Collen et al., 2013) | 3 |
| 19 | Mountain Green Cover Index | (FAO, 2017) | 1 |
| 20 | Number of certified forest areas under sustainable management with verified impacts on biodiversity conservation | (Kraxner et al., 2017; Maesano et al., 2018) | 1 |
| 21 | Number of invasive alien species in national lists as per the Global Register of Introduced and Invasive Species | (Pagad, 2018) | 1 |
| 22 | Number of invasive alien species introduction events | (Pagad, 2018) | 1 |
| 23 | Percent of total land area that is under cultivation | (Robertson, 2009) | 1 |
| 24 | Proportion of land that is degraded over total land area | (UNCCD, 2022) | 1 |
| 25 | Proportion of terrestrial, freshwater and marine ecological regions which are conserved by protected areas or other effective area-based conservation measures | (Chen and Wu, 2019; UNSTATS, 2020) | 1 |
| 26 | Protected Area Connectedness Index (PARC-Connectedness) | (Santini et al., 2016) | 2 |
| 27 | Protected Area Isolation Index (PAI) | (Brennan et al., 2022) | 1 |
| 28 | Protected Connected (Protconn) index | (Saura et al., 2017) | 2 |
| 29 | Rate of invasive alien species spread | (Seebens et al., 2020) | 1 |
| 30 | Red List Index (wild relatives of domesticated animals, for utilized species, pollinating species, for internationally traded species, impacts of invasive alien species) | (Butchart et al., 2007) | 9 |
| 31 | Red List of Ecosystems | (Keith et al., 2015) | 5 |
| 32 | Species habitat Index | (Powers and Jetz, 2019) | 2 |
| 33 | Species Protection Index | (GEO BON, 2015) | 3 |
| 34 | Species richness/Changes in local terrestrial diversity (PREDICTS) | (Newbold et al., 2015) | 1 |
| 35 | Tree cover loss | (Hansen et al., 2013) | 1 |
| 36 | Wild bird index | (Sheehan et al., 2010) | 1 |

Table S3.3: Biodiversity indicators from the Kunming-Montreal global biodiversity framework (CBD, 2022) with focus on tracking the status of terrestrial biodiversity, but which were not included in our analysis because they have too high uncertainties regarding information on their workflow.

| Indicator name | Uncertainty description |
| --- | --- |
| Biomass of selected natural ecosystems | No indicator description with this specific name could be found. The indicator mentioned under ID A.0.2 also changes between the zero draft (CBD, 2020) and the 1^st^ draft (CBD, 2021) from the Living Planet Index (Collen et al., 2013) to the Species Habitat Index (GEO BON, 2015). |
| CMS Connectivity Indicator | Discussions on the exact formulation of this indicator were still ongoing in October 2022 (<https://www.cbd.int/doc/notifications/2022/ntf-2022-063-indicators-en.pdf> |
| Extent of natural ecosystems | According to CBD website (<https://www.post-2020indicators.org/>, updated in December 2022) this indicator is still in development |
| Extent of natural ecosystems by type | According to CBD website (<https://www.post-2020indicators.org/>, updated in December 2022) this indicator is still in development |
| Forest distribution | No indicator with this name could be found. While quite a few remote sensing data products exist which provide a forest distribution layer, no specific workflow or data sources could be identified that match with this indicator name. |
| Forest Fragmentation Index | Only a documentation (<https://clear.uconn.edu/publications/research/tech_papers/Hurd_et_al_ASPRS2002.pdf>) of a presentation given in 2002 could be found which reports the development of the 'Forest Fragmentation Index' and its application to a small study area. |
| Grassland and savannah extent | No information on a specific indicator with this name could be found. The habitat ‘grassland and savannah’ is defined by IPBES as 'units of analysis' (<https://zenodo.org/record/3975694#.YfgKcPjvLRY>), but no temporal trends seem to be currently available and a specific workflow description is lacking. |
| Increase in secondary natural forest cover | No reference could be found for this this indicator. The name of the indicator is a description which implies an intersection between the Global Forest change product and a live cover fraction product on secondary forest). However, no workflow description could be found to analyse which data sources are used. |
| Intact wilderness | No indicator with this specific name could be found. The term wilderness appears in several papers (e.g. (Dinerstein et al., 2020) in the context of using Human Footprint maps (Venter et al., 2016a, 2016b). However, this is not a description of an indicator that measures intactness over time, and it therefore remains unclear which specific indicator is meant. |
| Peatland extent and condition | The ‘peatland extent and condition' indicator is sometimes mentioned, but no specific information on a workflow and the underlying data sources could be found. |
| Percent of land and seas covered by biodiversity-inclusive spatial plans | According to the CBD indicator website (<https://www.post-2020indicators.org/>, updated in December 2022 ), this indicator is still in development. |
| Percentage of cropped landscapes with at least 10% natural land | No information on this indicator could be found. While quite a few remote sensing data products exist on crop lands, no specific workflow or data sources could be identified that match with this description. |
| Percentage of threatened species that are improving in status according to the Red List | No specific indicator with this name could be identified. While it is clear from the description of the indicator that the Red-List status is to be used, no specific workflow describing this procedure could be identified. |
| Progress towards sustainable forest management (10.2) | According to CBD indicator website (<https://www.post-2020indicators.org/>, updated in December 2022) the description of this workflow is pending. |
| Proportion of agricultural area under productive and sustainable agriculture (10.1) | According to CBD indicator website (<https://www.post-2020indicators.org/>, updated in December 2022) the description of this workflow is pending. |
| Rate of invasive alien species establishment | According to CBD website (<https://www.post-2020indicators.org/>, updated in December 2022) the description of this workflow is pending. |
| Rate of invasive species impact and rate of impact | According to CBD indicator website (<https://www.post-2020indicators.org/>, updated in December 2022) the description of this workflow is pending. |
| Status of Key Biodiversity Areas | According to CBD indicator website (<https://www.post-2020indicators.org/>, updated in December 2022) the description of this workflow is pending. |
| Species Status Index | No specific indicator with this name could be identified. However, it is clear from the indicator name that data on species distribution and/or population size data are relevant for this indicator. |
| Total climate regulation services provided by ecosystems by ecosystem type (System of Environmental Economic Accounts) | According to the CBD website (<https://www.post-2020indicators.org/>, updated in December 2022) both the description of this workflow is pending. |

Table S3.4: Classification of keywords of insufficiently documented biodiversity indicators (in the Kunming-Montreal global biodiversity framework) into potential usage of Essential Biodiversity Variables (EBVs), used in the sensitivity analysis of the EBV relevance. The EBV classification (version of 30 January 2023) is derived from the Group on Earth Observations Biodiversity Observation Network (GEO BON) which distinguishes six EBV classes and twenty-one EBVs (https://geobon.org/ebvs/what-are-ebvs/).

| EBV class | EBV | Text descriptions (keywords) |
| --- | --- | --- |
| Genetic  composition | Genetic diversity | - |
|  | Genetic differentiation | - |
|  | Effective population size | - |
|  | Inbreeding | - |
| Species  populations | Species distributions | key biodiversity areas, pollinators, species spread/impact, threatened species |
|  | Species abundances | mean species abundance |
| Species  traits | Morphology | - |
|  | Physiology | - |
|  | Phenology | - |
|  | Movement | - |
|  | Reproduction | - |
| Community composition | Community abundance | - |
|  | Taxonomic / phylogenetic diversity | - |
|  | Trait diversity | - |
|  | Interaction diversity | - |
| Ecosystem  functioning | Primary productivity | biomass, condition, services, status of area, invasive species impact, services |
|  | Ecosystem phenology | condition, status of area, invasive species impact, services |
|  | Ecosystem disturbance | fragmentation, degraded, invasive species spread |
| Ecosystem  structure | Ecosystem vertical profile | intact wilderness |
|  | Live cover fraction | extent, cover, intact wilderness, proportion/ percentage of area/ecosystem |
|  | Ecosystem distribution | distribution, ecosystem type, extent, list of areas, intact wilderness, proportion/ percentage of area/ecosystem, connectivity |

Figure S3.1 Usage of essential biodiversity variables (EBVs, upper axis) by biodiversity indicators (y-axis) that were not eligible for inclusion in the main analysis. The indicators on the y-axis focus on tracking the state of terrestrial biodiversity but have high uncertainties regarding their workflow (e.g. they are in development and/or have not publicly available information on their workflow). By using keywords in the description for each indicator (see Table S2.3 in Supplementary Material S2), we identified which EBVs are most likely relevant for that indicator. The statistics below the table provide the summary of this potential usage, the current usage by sufficiently described indicators (see Figure 3 in the main text), and the combined usage of both. In addition, scores for the combined and current usage are provided (assessed according to criteria in Table 2 of the main text) to highlight the difference in ranking when including the potential usage of EBVs from insufficiently documented biodiversity indicators. Combining these (46) potential usages with the (126) current usages of the data products in the documented indicators only improved relevance scores of products associated with the ecosystem phenology EBV (from 3 = poor to 2 = moderate). Other results remained similar to the analysis with only the documented indicators.

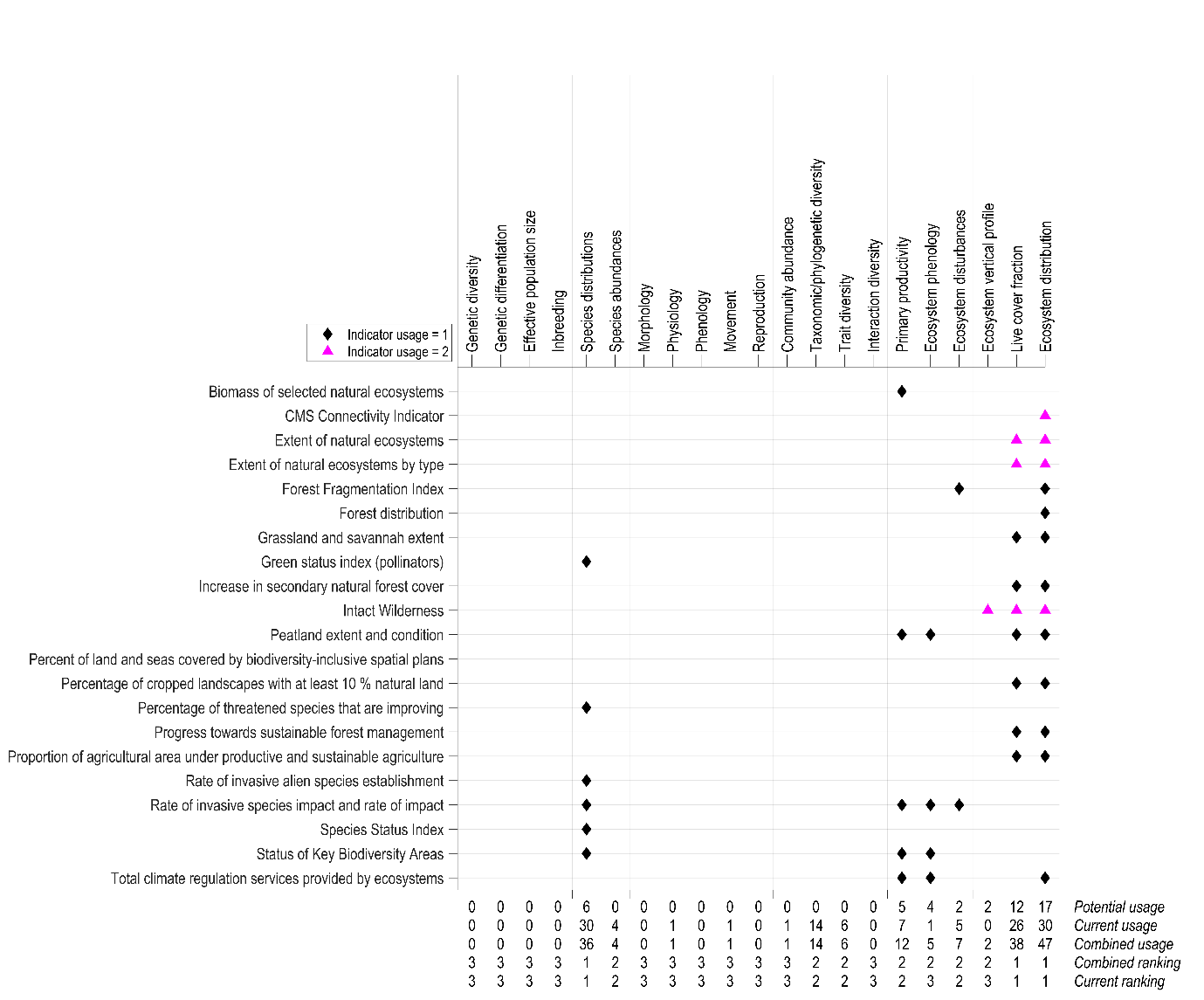
