## Supplementary Materials S4 for "Advancing terrestrial biodiversity monitoring with satellite remote sensing in the context of the Kunming-Montreal global biodiversity framework"

### Supplementary Material S4

Table S4.1: Analysis of spatial information products underlying the workflows of 35 biodiversity indicators (see Supplementary Material S3). These indicators are associated with the Kunming-Montreal global biodiversity framework (CBD, 2022) for tracking the status of terrestrial biodiversity. Each spatial information product was classified into the Essential Biodiversity Variable (EBV) framework (<https://geobon.org/ebvs/what-are-ebvs/>; accessed on 29 January 2023). For some spatial information products (such as threatened species), the data sources are only internally available within the responsible organization and consequently no data source could be provided here. Abbreviations are provided in Table S4.2. The indicator number refers to the biodiversity indicator as listed in Table S3.2 in Supplementary Material S3.

| ID | Variable name (as used in the biodiversity indicator workflows) | Indicator number | EBV classification | Spatial information products | Citation / responsible organisation |
| --- | --- | --- | --- | --- | --- |
| 1 | Above ground biomass | 1 | Primary productivity | SDG 15.2.1 | (FAO, 2020) |
| 2 | Agricultural land cover | 23 | Ecosystem distribution | Data from national reports | FAO |
| 3 | Alien Amphibians and reptiles | 29 | Species distributions | AmphRep | (Capinha et al., 2017) |
| 4 | Alien birds | 29 | Species distributions | GAVIA | (Dyer et al., 2017) |
| 5 | Alien invasive species | 29 | Species distributions | GRIIS | (Pagad, 2018) |
| 6 | Alien vascular plants | 29 | Species distributions | GloNAF | (van Kleunen et al., 2019) |
| 7 |  |  |  |  |  |
| 8 | Animal movement | 27 | Movement | Movebank | (Dodge et al., 2013) |
| 9 | Area of occupancy | 31 | Ecosystem distribution | Data by assessor | (Bland, L.M. Keith, D.A. , Miller and Murray, N.J., Rodríguez, 2017) |
| 10 | Beta diversity | 7,26 | Taxonomic/phylogenetic diversity | PREDICTS database | (Hoskins et al., 2020) |
| 11 | Beta diversity | 17 | Taxonomic/phylogenetic diversity | Global Safety Net layers | (Dinerstein et al., 2020) |
| 12 | Biodiversity hotspots | 17 | Species distributions | Global Safety Net layers | (Dinerstein et al., 2020) |
| 13 | Bird atlas information and distribution maps | 10 | Species distribution | Bird-Life International | (Peterjohn, 2006) |
| 14 | Bird population | 36 | Species distributions | Data from national surveys (Common Birds Census, the Breeding Bird Survey, the Wetland Bird Survey) | BTO, USGS, JNCC, RSPB, WWT |
| 15 | Bird Species abundance | 10 | Species abundance | PECBMS | (Sauer and Link, 2011) |
| 16 | Bird Species abundance | 10 | Species abundance | North American BBS | (Brlík et al., 2021) |
| 17 | Body weight | 27 | Physiology | PanTHERIA | (Jones et al., 2009) |
| 18 | Carbon stocks above and below ground | 24 | Primary productivity | HWSD | ISRIC |
| 19 | Carbon Stocks above and below ground | 24 | Primary productivity | GLOSIS | GSP |
| 20 | Conservation agriculture on arable land | 2 | Live cover fraction | Data from national surveys | FAO |
| 21 | Crop Area harvested | 2 | Ecosystem distribution | Data from national surveys | FAO |
| 22 | Crop wild relatives | 2 | Species distributions | Global distribution of CWR | (Crop Wild Relatives Occurrence data consortia, 2018) |
| 23 | Degraded land | 24 | Ecosystem disturbance | GLC-Share | FAO |
| 24 | Dietary Guild (herbivores, carnivores, omnivores) | 27 | Morphology | dietary guild | (Brennan et al., 2022) |
| 25 | Ecoregions | 32 | Ecosystem distribution | SRTM & ASTER DEM data | (Abrams et al., 2020) |
| 26 | Ecoregions | 28 | Ecosystem distributions | TEWD | (Olson et al., 2001) |
| 27 | Ecosystem quality | 11, 17 | Live cover fraction | Human Footprint Index | (Venter et al., 2016) |
| 28 | Energy balance, land surface albedo | 15 | Primary productivity | MCD43 | (Schaaf, C., Wang, 2015) |
| 29 | Energy balance, land surface temperature | 15 | Primary productivity | MOD11 | (Wan, 1999) |
| 30 | Energy balance, land cover dynamics | 15 | Primary productivity | MCD12Q | (Sulla-Menashe and Friedl, 2018) |
| 31 | Energy balance vegetation indices | 15 | Primary productivity | MYD13A3 | (Didan, 2015) |
| 32 | Extent of occurrence | 31 | Ecosystem distribution | Data by assessor | (Bland, L.M. Keith, D.A. , Miller and Murray, N.J., Rodríguez, 2017) |
| 33 | Forest area | 4 | Live cover fraction | Data from the forest management certification | (FAO, 2020) |
| 34 | Forest areas under sustainable management | 20 | Live cover fraction | Data from the forest management certification | (FAO, 2020) |
| 35 | Forested area and characteristics | 3, 6, 7, 13, 34, 35 | Live cover fraction | Global Forest Change product | (Hansen et al., 2013) |
| 36 | Fractional cover | 32 | Live cover fraction | LUH2 - fractional cover product | (Hurtt et al., 2020) |
| 37 | Gridded cell covered by primary vegetation | 16 | Primary productivity | MODIS vegetation condition maps | (NESDIS NOAA, 2013) |
| 38 | Habitat intactness | 12 | Ecosystem distributions | Habitat intactness index | (Beyer et al., 2020) |
| 39 | Important bird & biodiversity areas | 25, | Ecosystem distribution | protected areas key biodiversity areas | (UNEP-WCMC, 2019) |
| 40 | Intact large mammal assemblages | 17 | Community abundance | Global Safety Net layers | (Morrison et al., 2007) |
| 41 | Introduced and invasive species | 21 | Ecosystem distribution | Data from GRIIS | (Pagad, 2018) |
| 42 | IUCN range rarity sites | 17 | Species distributions | Global Safety Net layers | (Hill et al., 2019) |
| 43 | IUCN threatened species sites | 17 | Species distributions | Global Safety Net layers | (Hill et al., 2019) |
| 44 | Key biodiversity areas | 17 | Ecosystem distribution | WDPA | Birdlife international |
| 45 | Land cover | 14, 19 | Live cover fraction | Landcover CCI | ESA (ESA, 2017) |
| 46 | Land cover | 33 | Ecosystem distribution | Landsat 5, Landsat 7 and Landsat 8 | NASA |
| 47 | Land cover | 7 | Ecosystem distribution | CLC product | MCD12Q (Tuanmu and Jetz, 2014) |
| 48 | Land cover change | 15 | Ecosystem distribution | Global Forest Change product | (Hansen et al., 2013) |
| 49 | Land productivity degradation | 15 | Ecosystem disturbances | MYD13A3 | (Didan, 2015) |
| 50 | Land Productivity Dynamics | 24 | Primary productivity | MODIS NDVI/EVI | (Didan, 2015) |
| 51 | Land Productivity Dynamics | 24 | Primary productivity | ESA VGT-S | (Dierckx et al., 2014) |
| 52 | Land Productivity Dynamics | 24 | Primary productivity | GEOV1 NDVI | Copernicus |
| 53 | Land use | 7, 32 | Ecosystem distribution | LUH2 - land use product | (Hurtt et al., 2020, 2011) |
| 54 | Land use | 8, 12, 35 | Ecosystem distribution | Internal data products | (Hoskins et al., 2016) |
| 55 | Land use intensity | 8, 12, 35 | Ecosystem distribution | Internal data products | (Newbold et al., 2016) |
| 56 | Land use / cover (change) | 27, 33 | Ecosystem distribution | MCD12Q1 | (Sulla-Menashe and Friedl, 2018) |
| 57 | Land use / cover (change) | 26 | Ecosystem distribution | Data from WDPA | UNEP-WCMC |
| 58 | Last of the wild in each ecoregion | 17 | Species distributions | Global Safety Net layers | (Plumptre et al., 2019) |
| 59 | Livestock occurrence | 2 | Species distributions | GLW | (Robinson et al., 2014) |
| 60 | Local terrestrial diversity | 8, 34,12 | Taxonomic/phylogenetic diversity | PREDICTS database | (Hudson et al., 2014) |
| 61 | Measured Net primary production | 12 | Primary productivity | MOD13A1 | (Running and Zhao, 2015) |
| 62 | Mountain location | 19 | Ecosystem distribution | SRTM DEM | (Sayre et al., 2020) |
| 63 | Multi-taxonomic alien species | 29 | Species distributions | Alien Species FirstRecords | (Seebens et al., 2018) |
| 64 | Natural (biological) hazards | 16 | Ecosystem disturbances | Fire and Drought Risk maps | (NESDIS NOAA, 2013) |
| 65 | Number of Alien Invasive Species | 22 | Species distribution | GRIIS | (Pagad, 2018) |
| 66 | Open public (green/blue) Spaces |  | Ecosystem distribution | Open street map, Surveys, Google Earth Imagery | (UN-Habitat, 2018) |
| 67 | Organic agriculture on arable land | 2 | Live cover fraction | Data from national surveys | FAO |
| 68 | Pasture and cropland extents | 2 | Ecosystem distribution | Pasture and cropland extents | (Ramankutty et al., 2008) |
| 69 | Percentage of diversified cropland | 2 | Live cover fraction | SPAM crop physical areas | (Jones et al., 2021) |
| 70 | Percentage of cropland (>10% semi-natural vegetation) | 2 | Live cover fraction | Landcover CCI | (ESA, 2017) |
| 71 | Population | 33 | Species distributions | Rate of Decline, Range & Occurrence data | MOL |
| 72 | Population density | 8 | Live cover fraction | Data from GRUMP | (NASA and SECAC, 2016) |
| 73 | Population (Size/ density/ abundance/...) | 18 | Species abundance | Living Planet Index | (Collen et al., 2013) |
| 74 | Potential connectivity | 14 | Ecosystem distribution | Connectivity data | (Laestadius et al., 2011) |
| 75 | Effects of –agriculture on biodiversity | 14 | Ecosystem disturbances | Data on agriculture | USGS |
| 76 | Effects of deforestation on biodiversity | 14 | Ecosystem disturbances | Global Forest Change product | (Hansen et al., 2013) |
| 77 | Rare plant species | 17 | Species distributions | Global Safety Net layers | (Enquist et al., 2019) |
| 78 | Relative abundance of species functional types | 31 | Trait Diversity | Data by assessor | (Bland, L.M. Keith, D.A. , Miller and Murray, N.J., Rodríguez, 2017) |
| 79 | Relative abundance of guilds or alien species | 31 | Trait Diversity | Data by assessor | (Bland, L.M. Keith, D.A. , Miller and Murray, N.J., Rodríguez, 2017) |
| 80 | Small-range vertebrate sites | 17 | Species distributions | Global Safety Net layers | (Pimm et al., 2018) |
| 81 | Soil composition | 9 | Primary productivity | GSOC Map | (FAO and ITPS, 2020) |
| 82 | Soil Emission potentials | 9 | Primary productivity | WISE30Sec grids | ISRIC (Batjes, 2016) |
| 83 | Soil types | 9 | Primary productivity | Harmonized World Soil Database | FAO |
| 84 | Species composition | 31 | Taxonomic/phylogenetic diversity | Data by assessor | (Bland, L.M. Keith, D.A. , Miller and Murray, N.J., Rodríguez, 2017) |
| 85 | Species dominance | 31 | Taxonomic/phylogenetic diversity | Data by assessor | (Bland, L.M. Keith, D.A. , Miller and Murray, N.J., Rodríguez, 2017) |
| 86 | Species geographic distributions | 32 | Species distributions | MOL/GBIF records | MOL (Map of Life) |
| 87 | Species occurrence | 6, 7, 26 | Species distributions | MOL/GBIF records | (Hoskins et al., 2020) |
| 88 | Species richness | 31 | Taxonomic/phylogenetic diversity | Data by assessor | (Bland, L.M. Keith, D.A. , Miller and Murray, N.J., Rodríguez, 2017) |
| 89 | Temperature condition | 16 | Primary productivity | Brightness temperature (…) | (NESDIS NOAA, 2013) |
| 90 | Threatened species | 25, 30 | Species distributions | Red List | (Butchart et al., 2007) |
| 91 | Tropical primary forest land cover | 3 | Ecosystem distribution | Tropical primary forest land cover | (Turubanova et al., 2018) |
| 92 | Vegetation cover | 6, 26 | Live cover fraction | MODIS vegetation continuous fields | MOD44B (DiMiceli et al., 2011) |
| 93 | Vegetation cover | 27 | Live cover fraction | MYD13A3 | (Didan, 2015) |
| 94 | Vegetation coverage | 16 | Live cover fraction | Vegetation Health maps | (NESDIS NOAA, 2013) |
| 95 | Vegetation greenness | 16 | Ecosystem phenology | no noise NDVI maps | (NESDIS NOAA, 2013) |

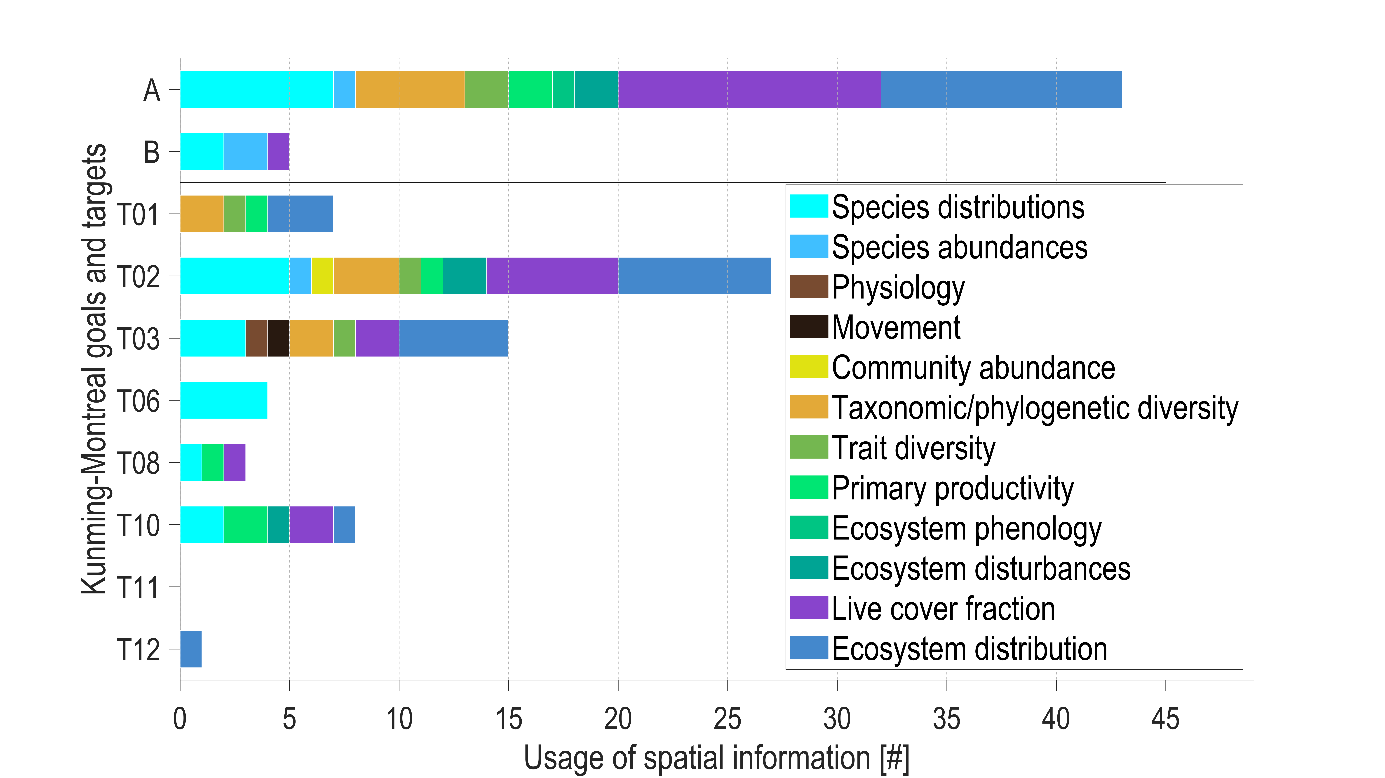

Figure S4.1: Usage of spatial information products related to essential biodiversity variables within the goals and targets of the Kunming-Montreal global biodiversity framework (CBD, 2022).

Table S4.2: Nomenclature for abbreviations in Table S4.1.

| Abbreviation | Description |
| --- | --- |
| ASTER | Advanced spaceborne thermal emission and reflection radiometer |
| BBS | Breeding Bird Survey |
| BILBI | Biogeographic modelling infrastructure for large-scale biodiversity indicators |
| BIP | Biodiversity indicator partnership |
| BTO | British trust for ornithology |
| CCI | Climate change initiative |
| CLC | Consensus land cover |
| CIESIN | [Center for international earth science information network](https://www.google.com/url?sa=t&rct=j&q=&esrc=s&source=web&cd=&ved=2ahUKEwi68IaiyLj2AhUlu6QKHXDkCHoQFnoECAgQAQ&url=http%3A%2F%2Fwww.ciesin.org%2F&usg=AOvVaw3eGRCQwC6zCZUN7RCBYoJH) |
| CWR | Crop Wild Relatives |
| DEM | Digital elevation model |
| ESA | European space agency |
| FAO | Food and agriculture organisation of the United Nations |
| GBIF | Global biodiversity information facility |
| GLW | Gridded Livestock of the World |
| GLOSIS | Global Soil Information System |
| GSOC | Global soil organic carbon |
| GRIIS | Global register of introduced and invasive species |
| GRUMP | Global rural-urban mapping project |
| HWSD | Harmonized World Soil Database |
| ISRIC | International soil reference and information centre |
| ITPS | Intergovernmental technical panel on soils |
| IUCN | International union for conservation of nature |
| JNCC | Joint nature conservation committee |
| LUH2 | Land use harmonisation |
| MODIS | Moderate resolution imaging spectroradiometer |
| MOL | Map of life |
| NDVI | Normalized difference vegetation index |
| PECBMS | Pan-European Common Birds Monitoring Scheme |
| PREDICTS | Projecting responses of ecological diversity in changing terrestrial systems |
| RSPB | Royal society for the protection of birds |
| SPAM | Spatial Production Allocation Model |
| SDG | Sustainable development goal |
| SRTM | Shuttle radar topography mission |
| UN | United nations |
| UNEP | United nations environmental programme |
| USGS | United States geological survey |
| WCMC | World conservation monitoring centre |
| WDKBA | World database on protected areas key biodiversity |
| WDPA | World database on protected areas |
| WISE30sec | World soil property estimates for broad- scale modelling |
| WWT | Wildfowl and wetland trust |

Hudson, L.N., Newbold, T., Contu, S., Hill, S.L.L., Lysenko, I., De Palma, A., Phillips, H.R.P., Senior, R.A., Bennett, D.J., Booth, H., Choimes, A., Correia, D.L.P., Day, J., Echeverría-Londoño, S., Garon, M., Harrison, M.L.K., Ingram, D.J., Jung, M., Kemp, V., Kirkpatrick, L., Martin, C.D., Pan, Y., White, H.J., Aben, J., Abrahamczyk, S., Adum, G.B., Aguilar-Barquero, V., Aizen, M.A., Ancrenaz, M., Arbeláez-Cortés, E., Armbrecht, I., Azhar, B., Azpiroz, A.B., Baeten, L., Báldi, A., Banks, J.E., Barlow, J., Batáry, P., Bates, A.J., Bayne, E.M., Beja, P., Berg, Å., Berry, N.J., Bicknell, J.E., Bihn, J.H., Böhning-Gaese, K., Boekhout, T., Boutin, C., Bouyer, J., Brearley, F.Q., Brito, I., Brunet, J., Buczkowski, G., Buscardo, E., Cabra-García, J., Calviño-Cancela, M., Cameron, S.A., Cancello, E.M., Carrijo, T.F., Carvalho, A.L., Castro, H., Castro-Luna, A.A., Cerda, R., Cerezo, A., Chauvat, M., Clarke, F.M., Cleary, D.F.R., Connop, S.P., D’Aniello, B., da Silva, P.G., Darvill, B., Dauber, J., Dejean, A., Diekötter, T., Dominguez-Haydar, Y., Dormann, C.F., Dumont, B., Dures, S.G., Dynesius, M., Edenius, L., Elek, Z., Entling, M.H., Farwig, N., Fayle, T.M., Felicioli, A., Felton, A.M., Ficetola, G.F., Filgueiras, B.K.C., Fonte, S.J., Fraser, L.H., Fukuda, D., Furlani, D., Ganzhorn, J.U., Garden, J.G., Gheler-Costa, C., Giordani, P., Giordano, S., Gottschalk, M.S., Goulson, D., Gove, A.D., Grogan, J., Hanley, M.E., Hanson, T., Hashim, N.R., Hawes, J.E., Hébert, C., Helden, A.J., Henden, J.A., Hernández, L., Herzog, F., Higuera-Diaz, D., Hilje, B., Horgan, F.G., Horváth, R., Hylander, K., Isaacs-Cubides, P., Ishitani, M., Jacobs, C.T., Jaramillo, V.J., Jauker, B., Jonsell, M., Jung, T.S., Kapoor, V., Kati, V., Katovai, E., Kessler, M., Knop, E., Kolb, A., Korösi, Á., Lachat, T., Lantschner, V., Le Féon, V., Lebuhn, G., Légaré, J.P., Letcher, S.G., Littlewood, N.A., López-Quintero, C.A., Louhaichi, M., Lövei, G.L., Lucas-Borja, M.E., Luja, V.H., Maeto, K., Magura, T., Mallari, N.A., Marin-Spiotta, E., Marshall, E.J.P., Martínez, E., Mayfield, M.M., Mikusinski, G., Milder, J.C., Miller, J.R., Morales, C.L., Muchane, M.N., Muchane, M., Naidoo, R., Nakamura, A., Naoe, S., Nates-Parra, G., Navarrete Gutierrez, D.A., Neuschulz, E.L., Noreika, N., Norfolk, O., Noriega, J.A., Nöske, N.M., O’Dea, N., Oduro, W., Ofori-Boateng, C., Oke, C.O., Osgathorpe, L.M., Paritsis, J., Parra-H, A., Pelegrin, N., Peres, C.A., Persson, A.S., Petanidou, T., Phalan, B., Philips, T.K., Poveda, K., Power, E.F., Presley, S.J., Proença, V., Quaranta, M., Quintero, C., Redpath-Downing, N.A., Reid, J.L., Reis, Y.T., Ribeiro, D.B., Richardson, B.A., Richardson, M.J., Robles, C.A., Römbke, J., Romero-Duque, L.P., Rosselli, L., Rossiter, S.J., Roulston, T.H., Rousseau, L., Sadler, J.P., Sáfián, S., Saldaña-Vázquez, R.A., Samnegård, U., Schüepp, C., Schweiger, O., Sedlock, J.L., Shahabuddin, G., Sheil, D., Silva, F.A.B., Slade, E.M., Smith-Pardo, A.H., Sodhi, N.S., Somarriba, E.J., Sosa, R.A., Stout, J.C., Struebig, M.J., Sung, Y.H., Threlfall, C.G., Tonietto, R., Tóthmérész, B., Tscharntke, T., Turner, E.C., Tylianakis, J.M., Vanbergen, A.J., Vassilev, K., Verboven, H.A.F., Vergara, C.H., Vergara, P.M., Verhulst, J., Walker, T.R., Wang, Y., Watling, J.I., Wells, K., Williams, C.D., Willig, M.R., Woinarski, J.C.Z., Wolf, J.H.D., Woodcock, B.A., Yu, D.W., Zaitsev, A.S., Collen, B., Ewers, R.M., Mace, G.M., Purves, D.W., Scharlemann, J.P.W., Purvis, A., 2014. The PREDICTS database: A global database of how local terrestrial biodiversity responds to human impacts. Ecology and Evolution 4, 4701–4735. https://doi.org/10.1002/ece3.1303

Hurtt, G.C., Chini, L., Sahajpal, R., Frolking, S., Bodirsky, B.L., Calvin, K., Doelman, J.C., Fisk, J., Fujimori, S., Goldewijk, K.K., Hasegawa, T., Havlik, P., Heinimann, A., Humpenöder, F., Jungclaus, J., Kaplan, J.O., Kennedy, J., Krisztin, T., Lawrence, D., Lawrence, P., Ma, L., Mertz, O., Pongratz, J., Popp, A., Poulter, B., Riahi, K., Shevliakova, E., Stehfest, E., Thornton, P., Tubiello, F.N., van Vuuren, D.P., Zhang, X., 2020. Harmonization of global land use change and management for the period 850-2100 (LUH2) for CMIP6. Geoscientific Model Development 13, 5425–5464. https://doi.org/10.5194/gmd-13-5425-2020

Hurtt, G.C., Chini, L.P., Frolking, S., Betts, R.A., Feddema, J., Fischer, G., Fisk, J.P., Hibbard, K., Houghton, R.A., Janetos, A., Jones, C.D., Kindermann, G., Kinoshita, T., Klein Goldewijk, K., Riahi, K., Shevliakova, E., Smith, S., Stehfest, E., Thomson, A., Thornton, P., van Vuuren, D.P., Wang, Y.P., 2011. Harmonization of land-use scenarios for the period 1500-2100: 600 years of global gridded annual land-use transitions, wood harvest, and resulting secondary lands. Climatic Change 109, 117–161. https://doi.org/10.1007/s10584-011-0153-2

Jones, K.E., Bielby, J., Cardillo, M., Fritz, S.A., O’Dell, J., Orme, C.D.L., Safi, K., Sechrest, W., Boakes, E.H., Carbone, C., Connolly, C., Cutts, M.J., Foster, J.K., Grenyer, R., Habib, M., Plaster, C.A., Price, S.A., Rigby, E.A., Rist, J., Teacher, A., Bininda-Emonds, O.R.P., Gittleman, J.L., Mace, G.M., Purvis, A., 2009. PanTHERIA: a species‐level database of life history, ecology, and geography of extant and recently extinct mammals. Ecology 90, 2648–2648. https://doi.org/10.1890/08-1494.1

Jones, S.K., Estrada-Carmona, N., Juventia, S.D., Dulloo, M.E., Laporte, M.A., Villani, C., Remans, R., 2021. Agrobiodiversity Index scores show agrobiodiversity is underutilized in national food systems. Nature Food 2, 712–723. https://doi.org/10.1038/s43016-021-00344-3

Laestadius, L., Maginnis, S., Minnemeyer, S., Potapov, P., Saint-Laurent, C., Sizer, N., 2011. Mapping opportunities for forest landscape restoration. Unasylva 62, 47–48.

Morrison, J.C., Sechrest, W., Dinerstein, E., Wilcove, D.S., Lamoreux, J.F., 2007. Persistence of large mammal faunas as indicators of global human impacts. Journal of Mammalogy 88, 1363–1380. https://doi.org/10.1644/06-MAMM-A-124R2.1

NASA, SECAC, 2016. Gridded Population of the World (GPW), v4. Center for International Earth Science Information Network - CIESIN - Columbia University. https://doi.org/10.7927/H4F47M2C

NESDIS NOAA, 2013. AVHRR Vegetation Health Product (AVHRR-VHP) User Guide [WWW Document]. URL www.star.nesdis.noaa.gov/smcd/emb/vci/VH_doc/VHP_uguide_v1.4_2013_1221.pdf (assessed 1 May 2022)

Newbold, T., Hudson, L.N., Arnell, A.P., Contu, S., De Palma, A., Ferrier, S., Hill, S.L.L., Hoskins, A.J., Lysenko, I., Phillips, H.R.P., Burton, V.J., Chng, C.W.T., Emerson, S., Gao, D., Hale, G.P., Hutton, J., Jung, M., Sanchez-Ortiz, K., Simmons, B.I., Whitmee, S., Zhang, H., Scharlemann, J.P.W., Purvis, A., 2016. Has land use pushed terrestrial biodiversity beyond the planetary boundary? A global assessment. Science 353, 291–288. https://doi.org/10.1126/science.aaf2201

Olson, D.M., Dinerstein, E., Wikramanayake, E.D., Burgess, N.D., Powell, G.V.N., Underwood, E.C., D’Amico, J.A., Itoua, I., Strand, H.E., Morrison, J.C., Loucks, C.J., Allnutt, T.F., Ricketts, T.H., Kura, Y., Lamoreux, J.F., Wettengel, W.W., Hedao, P., Kassem, K.R., 2001. Terrestrial ecoregions of the world: A new map of life on Earth. BioScience 51, 933–938. https://doi.org/10.1641/0006-3568(2001)051[0933:TEOTWA]2.0.CO;2

Pagad, S., 2018. Introducing the Global Register of Introduced and Invasive Species: challenges with classification. Biodiversity Information Science and Standards 2, e25306. https://doi.org/10.3897/biss.2.25306

Peterjohn, B., 2006. Birds in Europe: Population Estimates, Trends and Conservation Status. The Auk 123, 915–916. https://doi.org/10.1093/auk/123.3.915

Pimm, S.L., Jenkins, C.N., Li, B. V., 2018. How to protect half of earth to ensure it protects sufficient biodiversity. Science Advances 4, eaat2616. https://doi.org/10.1126/sciadv.aat2616

Plumptre, A.J., Baisero, D., Jędrzejewski, W., Kühl, H., Maisels, F., Ray, J.C., Sanderson, E.W., Strindberg, S., Voigt, M., Wich, S., 2019. Are We Capturing Faunal Intactness? A Comparison of Intact Forest Landscapes and the “Last of the Wild in Each Ecoregion.” Frontiers in Forests and Global Change 2, 24. https://doi.org/10.3389/ffgc.2019.00024

Ramankutty, N., Evan, A.T., Monfreda, C., Foley, J.A., 2008. Farming the planet: 1. Geographic distribution of global agricultural lands in the year 2000. Global Biogeochemical Cycles 22, eGB1003. https://doi.org/10.1029/2007GB002952

Robinson, T.P., William Wint, G.R., Conchedda, G., Van Boeckel, T.P., Ercoli, V., Palamara, E., Cinardi, G., D’Aietti, L., Hay, S.I., Gilbert, M., 2014. Mapping the global distribution of livestock. PLoS ONE 9, e96084. https://doi.org/10.1371/journal.pone.0096084

Running, S.W., Zhao, M., 2015. User’s Guide Daily GPP and Annual NPP (MOD17A2/A3) Products, V3 ed. NASA Earth Observing System MODIS Land Algorithm.

Sauer, J.R., Link, W.A., 2011. Analysis of the North American Breeding Bird Survey using hierarchical models. Auk 128, 87–98. https://doi.org/10.1525/auk.2010.09220

Sayre, R., Karagulle, D., Frye, C., Boucher, T., Wolff, N.H., Breyer, S., Wright, D., Martin, M., Butler, K., Van Graafeiland, K., Touval, J., Sotomayor, L., McGowan, J., Game, E.T., Possingham, H., 2020. An assessment of the representation of ecosystems in global protected areas using new maps of World Climate Regions and World Ecosystems. Global Ecology and Conservation 21, e00860. https://doi.org/10.1016/j.gecco.2019.e00860

Schaaf, C., Wang, Z., 2015. MCD43A1 MODIS/Terra+Aqua BRDF/Albedo Model Parameters Daily L3 Global - 500m V006 [Data set]. NASA EOSDIS Land Processes DAAC. https://doi.org/10.5067/MODIS/MCD43A1.006

Seebens, H., Blackburn, T.M., Dyer, E.E., Genovesi, P., Hulme, P.E., Jeschke, J.M., Pagad, S., Pyšek, P., Van Kleunen, M., Winter, M., Ansong, M., Arianoutsou, M., Bacher, S., Blasius, B., Brockerhoff, E.G., Brundu, G., Capinha, C., Causton, C.E., Celesti-Grapow, L., Dawson, W., Dullinger, S., Economo, E.P., Fuentes, N., Guénard, B., Jäger, H., Kartesz, J., Kenis, M., Kühn, I., Lenzner, B., Liebhold, A.M., Mosena, A., Moser, D., Nentwig, W., Nishino, M., Pearman, D., Pergl, J., Rabitsch, W., Rojas-Sandoval, J., Roques, A., Rorke, S., Rossinelli, S., Roy, H.E., Scalera, R., Schindler, S., Štajerová, K., Tokarska-Guzik, B., Walker, K., Ward, D.F., Yamanaka, T., Essl, F., 2018. Global rise in emerging alien species results from increased accessibility of new source pools. Proceedings of the National Academy of Sciences of the United States of America 115, 2264–2273. https://doi.org/10.1073/pnas.1719429115

Sulla-Menashe, D., Friedl, M.A., 2018. User Guide to Collection 6 MODIS Land Cover Dynamics (MCD12Q2) Product, User Guide. https://doi.org/doi.org/10.5067/MODIS/MCD12Q1.006

Tuanmu, M.N., Jetz, W., 2014. A global 1-km consensus land-cover product for biodiversity and ecosystem modelling. Global Ecology and Biogeography 23, 1031–1045. https://doi.org/10.1111/geb.12182

Turubanova, S., Potapov, P. V., Tyukavina, A., Hansen, M.C., 2018. Ongoing primary forest loss in Brazil, Democratic Republic of the Congo, and Indonesia. Environmental Research Letters 13, e074028. https://doi.org/10.1088/1748-9326/aacd1c

UN-Habitat, 2018. SDG Indicator 11.7.1 Training Module: Public Space. United Nations Human Settlement Programme (UN-Habitat), Nairobi, https://unhabitat.org/sites/default/files/2020/07/indicator_11.7.1_training_module_public_space.pdf.

UNEP-WCMC, 2019. User Manual for the World Database on Protected Areas and world database on other effective area-based conservation measures: 1.6 [WWW Document]. NEP-WCMC: Cambridge, UK. URL http://wcmc.io/WDPA_Manual (accessed 1 May 2022)

van Kleunen, M., Pyšek, P., Dawson, W., Essl, F., Kreft, H., Pergl, J., Weigelt, P., Stein, A., Dullinger, S., König, C., Lenzner, B., Maurel, N., Moser, D., Seebens, H., Kartesz, J., Nishino, M., Aleksanyan, A., Ansong, M., Antonova, L.A., Barcelona, J.F., Breckle, S.W., Brundu, G., Cabezas, F.J., Cárdenas, D., Cárdenas-Toro, J., Castaño, N., Chacón, E., Chatelain, C., Conn, B., de Sá Dechoum, M., Dufour-Dror, J.M., Ebel, A.L., Figueiredo, E., Fragman-Sapir, O., Fuentes, N., Groom, Q.J., Henderson, L., Inderjit, Jogan, N., Krestov, P., Kupriyanov, A., Masciadri, S., Meerman, J., Morozova, O., Nickrent, D., Nowak, A., Patzelt, A., Pelser, P.B., Shu, W. sheng, Thomas, J., Uludag, A., Velayos, M., Verkhosina, A., Villaseñor, J.L., Weber, E., Wieringa, J.J., Yazlık, A., Zeddam, A., Zykova, E., Winter, M., 2019. The Global Naturalized Alien Flora (GloNAF) database. Ecology 100, e02542. https://doi.org/10.1002/ecy.2542

Venter, O., Sanderson, E.W., Magrach, A., Allan, J.R., Beher, J., Jones, K.R., Possingham, H.P., Laurance, W.F., Wood, P., Fekete, B.M., Levy, M.A., Watson, J.E.M., 2016. Sixteen years of change in the global terrestrial human footprint and implications for biodiversity conservation. Nature Communications 7, e12558. https://doi.org/10.1038/ncomms12558

Wan, Z., 1999. MODIS land-surface temperature algorithm theoretical basis document (LST ATBD). NASA, Washington DC, USA.
