## Supplementary Materials S1 for "Advancing terrestrial biodiversity monitoring with satellite remote sensing in the context of the Kunming-Montreal global biodiversity framework"

### Supplementary Material S5

Table S5.1: Ranking of products from satellite remote sensing (SRS) based on summary scores of the feasibility, accuracy, relevance and immaturity criteria (see main text for details on scoring). Each SRS product is classified into an Essential Biodiversity Variable (EBV). The top priority products (rank ≤ 7) are highlighted in light grey. Scores and ranks in brackets for denote their updated value if potential EBV usage (from biodiversity indicators that are currently insufficiently documented) are taken into account. Products highlighted with a star (*) are omitted from the ranking, as they do not report on the state of biodiversity, but instead of the response to disturbances.

| SRS products | EBV | Scoring | | | | | Rank |
| --- | --- | --- | --- | --- | --- | --- | --- |
|  |  | Feasibility | Accuracy | Relevance | Immaturity | Total |  |
| Fraction of vegetation cover | Live cover fraction | 1 | 2 | 1 | 2 | 6 | 1 |
| Plant area index profile (canopy cover) | Live cover fraction | 1 | 2 | 1 | 2 | 6 | 1 |
| Above-ground biomass | Ecosystem distribution | 1.5 | 1 | 2 | 2 | 6.5 | 3 |
| Foliar N/P/K content | Primary productivity | 1.5 | 1 | 2 | 2 | 6.5 | 3 |
| Land cover (Vegetation type) | Live cover fraction | 1 | 1.5 | 1 | 3 | 6.5 | 3 |
| Leaf area index | Physiology | 1 | 1.5 | 3 | 1 | 6.5 | 3 |
| Carbon cycle (above-ground biomass) | Primary productivity | 1 | 2 | 2 | 2 | 7 | 7 |
| Chlorophyll content and flux | Physiology | 1 | 2 | 3 | 1 | 7 | 7 |
| Ecosystem fragmentation | Ecosystem distribution | 1 | 2 | 2 | 2 | 7 | 7 |
| Ecosystem structural variance | Ecosystem distribution | 1 | 2 | 2 | 2 | 7 | 7 |
| Gross primary productivity | Physiology | 1 | 2 | 3 | 1 | 7 | 7 |
| Leaf area index | Ecosystem distribution | 1 | 1.5 | 2 | 3 | 7.5 | 12 |
| Leaf area index | Primary productivity | 1 | 1.5 | 2 | 3 | 7.5 | 12 |
| Carbon cycle (below ground biomass and carbon) | Primary productivity | 2 | 3 | 2 | 1 | 8 | 14 |
| Carbon cycle (sequestration) | Primary productivity | 2 | 3 | 2 | 1 | 8 | 14 |
| Chlorophyll content and flux | Primary productivity | 2 | 2 | 2 | 2 | 8 | 14 |
| Evapotranspiration | Primary productivity | 2 | 3 | 2 | 1 | 8 | 14 |
| Foliar N/P/K content | Physiology | 2 | 2 | 3 | 1 | 8 | 14 |
| Fraction of absorbed photosynthetically active radiation | Primary productivity | 1 | 2 | 2 | 3 | 8 | 14 |
| Functional diversity | Taxonomic/phylogenetic diversity | 2 | 3 | 2 | 1 | 8 | 14 |
| Green-up (start of season) | Phenology | 2 | 2 | 3 | 1 | 8 | 14 |
| Gross primary productivity | Primary productivity | 1 | 2 | 2 | 3 | 8 | 14 |
| Habitat structure | Ecosystem vertical profile | 1 | 2 | 3 | 2 | 8 | 14 |
| Ice cover habitat | Ecosystem vertical profile | 1 | 1 | 3 | 3 | 8 | 14 |
| Leaf dry matter content | Morphology | 2 | 2 | 3 | 1 | 8 | 14 |
| Net primary productivity | Physiology | 2 | 2 | 3 | 1 | 8 | 14 |
| Peak season (max of season) | Phenology | 2 | 2 | 3 | 1 | 8 | 14 |
| Population density (distribution) | Species abundances | 2 | 3 | 2 | 1 | 8 | 14 |
| Relative species abundance | Species abundances | 2 | 3 | 2 | 1 | 8 | 14 |
| Senescence (end of season) | Phenology | 2 | 2 | 3 | 1 | 8 | 14 |
| Species abundance | Species abundances | 2 | 3 | 2 | 1 | 8 | 14 |
| Species diversity indices (Simpson, Shannon, alpha, beta, gamma) | Species distributions | 3 | 3 | 1 | 1 | 8 | 14 |
| Species richness | Species distributions | 3 | 3 | 1 | 1 | 8 | 14 |
| Specific leaf area | Morphology | 2 | 2 | 3 | 1 | 8 | 14 |
| Specific leaf area | Primary productivity | 2 | 2 | 2 | 2 | 8 | 14 |
| Urban habitat | Ecosystem distribution | 1 | 2 | 2 | 3 | 8 | 14 |
| Vegetation canopy height | Ecosystem vertical profile | 1 | 2 | 3 | 2 | 8 | 14 |
| Cellulose | Physiology | 2 | 3 | 3 | 1 | 9 | 38 (41) |
| Ecosystem soil moisture | Primary productivity | 3 | 2 | 2 | 2 | 9 | 38 (41) |
| Forest species and age class | Species abundances | 3 | 3 | 2 | 1 | 9 | 38 (41) |
| Land surface phenology green-up (start of season) | Ecosystem phenology | 2 | 2 | 3 (2) | 2 | 9 (8) | 38 (14) |
| Land surface phenology peak (max of season) | Ecosystem phenology | 2 | 2 | 3 (2) | 2 | 9 (8) | 38 (14) |
| Land surface phenology senescence (end of season) | Ecosystem phenology | 2 | 2 | 3 (2) | 2 | 9 (8) | 38 (14) |
| Lignin | Physiology | 2 | 3 | 3 | 1 | 9 | 38 (41) |
| Net primary productivity | Primary productivity | 2 | 2 | 2 | 3 | 9 | 38 (41) |
| Non-structural carbohydrates | Physiology | 2 | 3 | 3 | 1 | 9 | 38 (41) |
| Phylogenetic diversity | Taxonomic/phylogenetic diversity | 3 | 3 | 2 | 1 | 9 | 38 (41) |
| Polyphenols | Physiology | 2 | 3 | 3 | 1 | 9 | 38 (41) |
| Taxonomic (Species diversity/ richness) | Taxonomic/phylogenetic diversity | 3 | 3 | 2 | 1 | 9 | 38 (41) |
| Deadwood habitat | Ecosystem vertical profile | 3 | 3 | 3 | 1 | 10 | 50 |
| Number or percentage of species which grow or occur together | Community abundance | 3 | 3 | 3 | 1 | 10 | 50 |
| Biological effects fire disturbance* | Ecosystem vertical profile | 1 | 1 | 3 | 3 | 8 | - |
| Biological effects fire disturbance* | Ecosystem disturbances | 1 | 1 | 2 | 3 | 8 | - |
| Biological effects of Irregular inundation* | Ecosystem vertical profile | 1 | 1 | 3 | 3 | 8 | - |
| Biological effects of Irregular inundation* | Ecosystem disturbances | 1 | 1 | 2 | 3 | 7 | - |
| Biological effects of Pest and disease outbreak* | Ecosystem disturbances | 2.5 | 2.5 | 2 | 1 | 8 | - |

Please note that the ranking prioritizes products with a high level of immaturity, but also with a high level of feasibility, accuracy and relevance so that they are interesting to be further developed. As such, this ranking prioritizes other products than those that are important for maintaining current missions by space agencies (Skidmore et al., 2021). Hence, the priority list above ranks immaturity high whereas the priority list from Skidmore et al. (2021) ranked maturity high (to highlight which current SRS product should be maintained). In comparison, we found that plant physiology products should be given a higher priority (see Figure S5.1) because SRS products of species traits (e.g. leaf area index, chlorophyll content and flux and gross primary productivity) are currently immature but can provide essential information for tracking biodiversity change, including plant phenology (e.g. timing of flowering and fruiting) and stoichiometry (e.g. the balance between carbon, nitrogen and phosphorus).

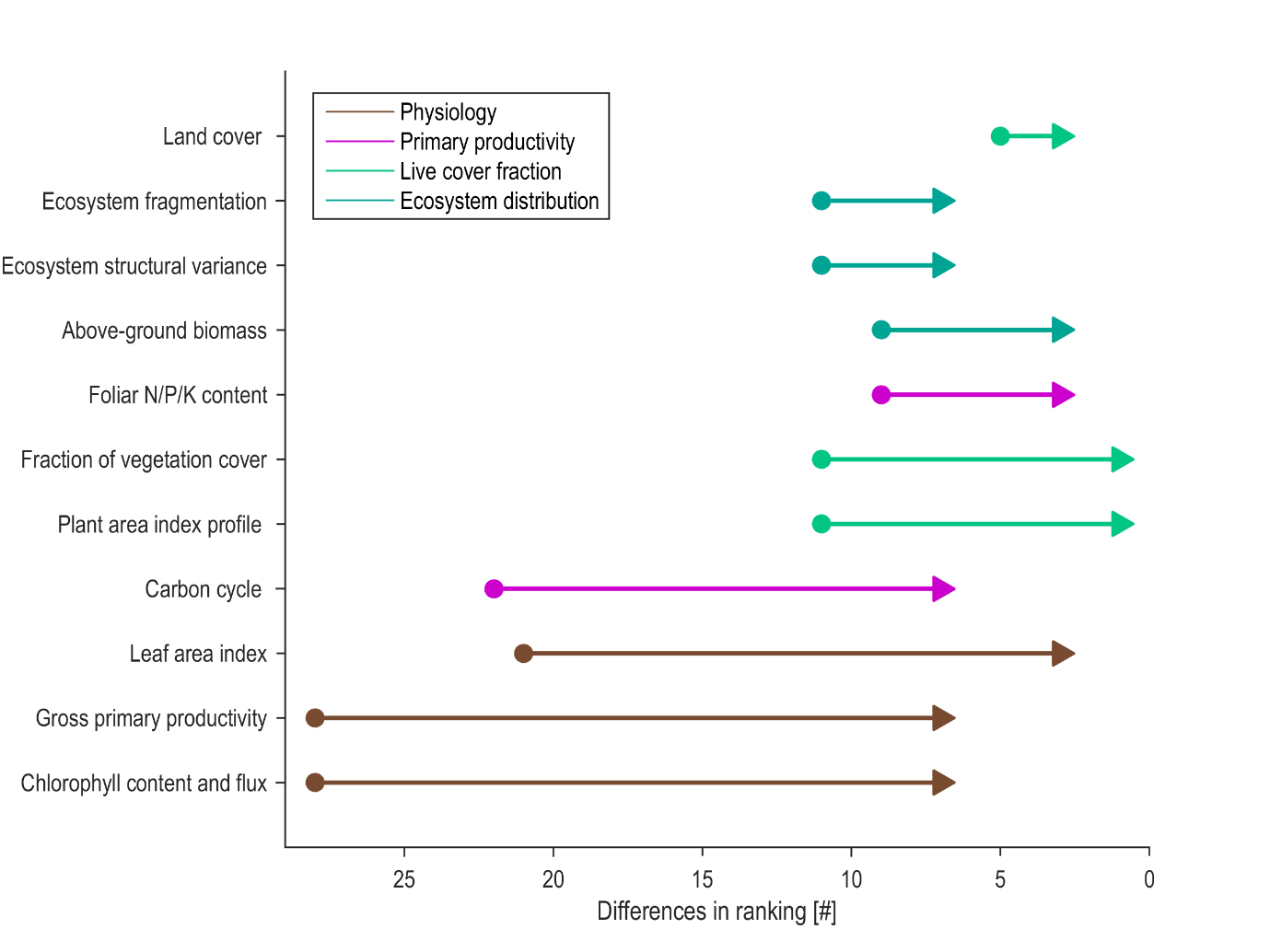

Figure S5.1. Changes in the ranking of satellite remote sensing products from the top priority list (Table 5, main text) when the development potential by the scientific community (our focus, represented by triangles) is compared to the operation and maintenance of missions by space agencies (focus of Skidmore et al. 2021, represented by closed circles). A lower rank (to the right) represents a higher priority.
